# Massively parallel protein-protein interaction measurement by sequencing (MP3-seq) enables rapid screening of protein heterodimers

**DOI:** 10.1101/2023.02.08.527770

**Authors:** Alexander Baryshev, Alyssa La Fleur, Benjamin Groves, Cirstyn Michel, David Baker, Ajasja Ljubetič, Georg Seelig

## Abstract

Protein-protein interactions (PPIs) regulate many cellular processes, and engineered PPIs have cell and gene therapy applications. Here we introduce massively parallel protein-protein interaction measurement by sequencing (MP3-seq), an easy-to-use and highly scalable yeast-two-hybrid approach for measuring PPIs. In MP3-seq, DNA barcodes are associated with specific protein pairs, and barcode enrichment can be read by sequencing to provide a direct measure of interaction strength. We show that MP3-seq is highly quantitative and scales to over 100,000 interactions. We apply MP3-seq to characterize interactions between families of rationally designed heterodimers and to investigate elements conferring specificity to coiled-coil interactions. Finally, we predict coiled heterodimer structures using AlphaFold-Multimer (AF-M) and train linear models on physics simulation energy terms to predict MP3-seq values. We find that AF-M and AF-M complex prediction-based models could be valuable for pre-screening interactions, but that measuring interactions experimentally remains necessary to rank their strengths quantitatively.

## Introduction

Synthetic protein binders that can mediate interactions between other proteins or cells have the potential to revolutionize fields from cell therapy (Cho et al., 2018) to synthetic biology (Chen et al., 2020; Chen and Elowitz, 2021; Gao et al., 2018; Groves et al., 2016) and material science (Ben-Sasson et al., 2021; Gonen et al., 2015; Ljubetič et al., 2017b). Early work in synthetic biology often relied on natural interaction domains such as SH3 and PDZ (Kuriyan and Cowburn, 1997). However, such components provide a poor starting point for rationally designing large-scale assemblies due to crosstalk. Consequently, synthetic protein circuits remain much smaller and simpler than biological PPI networks. In order to scale up synthetic protein-based circuits, we need large-scale libraries of modular interaction domains.

Fulfilling this need through rational heterodimer design has yielded promising results, but creating large, totally orthogonal sets of dimers remains challenging. Early rational design work focused on coiled-coil dimers (1×1s): 1×1 coil binding is primarily determined by complementary electrostatic and hydrophobic interactions at specific heptad positions. Because the biophysical rules guiding these interactions are relatively well understood and can be captured by predictive models (Fong et al., 2004; Potapov et al., 2015), orthogonal sets of up to six 1×1 heterodimers have been generated (Lebar et al., 2020; Thompson et al., 2012). Recently, a set of orthogonal 1×1s (primarily homodimers) was identified in a high-throughput screen (Boldridge et al., 2020). However, the restricted geometry required for 1×1 coil interactions limits the number of possible orthogonal interactions. The Baker lab expanded the coiled-coil toolbox by introducing helical bundle heterodimers, with protomers consisting of two helices connected by a hairpin loop, where interactions are determined by designed hydrogen bond networks between the bundles (we will refer to these as 2×2s) (Boyken et al., 2016; Chen et al., 2019). This multi-helix bundle approach increases the alphabet of possible coil interactions and could pave the way to generating larger orthogonal sets. However, reliably minimizing off-target interactions (e.g., due to proteins associating in an unexpected register or orientation) remains challenging as these are not captured in the biophysical design objective to produce 2×2s. A practical alternative is to prepare diverse libraries of *de novo* 2×2 protomers and experimentally measure an all-by-all matrix of interactions. Afterward, an orthogonal set can be extracted (Brodnik et al., 2019).

Many PPI measuring methods have been developed which could be used to run all-by-all PPI screens with varying throughput levels. Mass spectrometry and protein-array based methods can measure PPIs but require laborious protein purification steps (Rao et al., 2014; Smits and Vermeulen, 2016). Yeast- or phage-display methods use next-generation sequencing technologies to increase throughput but are limited to “several-versus-many” screening (Rouet et al., 2018). Alpha-seq, a recent high-throughput method that takes advantage of the yeast mating pathway, overcomes this limitation and allows library-on-library screening (Younger et al., 2017). However, throughput is still limited, and not all proteins correctly fold when displayed on the yeast surface.

Yeast two-hybrid (Y2H) methods are a powerful alternative to surface display for characterizing PPIs (Fields and Song, 1989). In Y2H, one protein of interest is fused to a DNA binding domain (DBD) and the second is fused to a transcriptional activation domain (AD). If the proteins interact, a functional transcription factor is reconstituted and drives the expression of a growth-essential enzyme. Early Y2H approaches tested small numbers of PPIs using plate-based selection, but lab automation and pooling strategies (Luck et al., 2020; Weile et al., 2017) have enabled proteome-scale screens. To further address Y2H scaling issues, high-throughput Y2H (HT-Y2H) and enzyme complementation methods have been developed leveraging next-generation sequencing to read out interaction strength (Diss and Lehner, 2018; Erffelinck et al., 2018; Jin et al., 2007; Rajagopala and Uetz, 2009; Trigg et al., 2017; Weimann et al., 2013; Yachie et al., 2016; F. Yang et al., 2018; J.-S. Yang et al., 2018; Yu et al., 2011). Most of these methods require library construction in f. *coli* before screening interactions in yeast or rely on yeast mating and thus need separate protein libraries to transform MATa and MATa yeast. High-throughput bacterial two-hybrid assays have been developed, which provide a non-eukaryotic alternative for screening PPIs (Boldridge et al., 2020). Concomitantly with experimental methods, custom workflows for analyzing HT-Y2H data were developed (Banerjee et al., 2021; Velasquez-Zapata et al., 2021).

Recently, deep learning models such as AlphaFold2 (Jumper et al., 2021) and RoseTTAFold (Baek et al., 2021) have been shown to predict protein structures with near experimental accuracy. Structure prediction for proteins with multiple chains is possible, either through the concatenation of target chains with a flexible linker, or with specialized models such as AlphaFold-Multimer (AF-M) (Jumper et al., 2021). AF-M takes in multiple input sequences and has been shown to improve interface predictions over AlphaFold2. Given the speed at which these models operate compared to running Y2H assays, even high throughput ones, determining how these models can be used to pre-screen PPIS or supplement PPI assays is highly attractive. Tools have been developed to run AF-M in an all-by-all manner to assist with PPI pre-screening (Yu et al., 2022), and AF-M error metrics have been demonstrated to be state-of-the-art protein-peptide interaction predictors (Johansson-Akhe and Wallner, 2022). However, it is unclear how accurately non-structural model derived *de novo* dimerization can be predicted with AF-M for structurally similar proteins and if AF-M predictions can be used to find orthogonal protein sets.

Here, we introduced MP3-seq, a massively parallel Y2H workflow for measuring PPIs using sequencing. In MP3-seq, the identity of each protein is encoded in a DNA barcode, and the relative barcode-barcode pair abundance before and after a selection experiment serves as a proxy for interaction strength. Plasmids are assembled in yeast through homologous recombination encoding the protein pairs of interest, their associated barcodes, and all other elements required for Y2H experiments. Thus, our experimental workflow bypasses the need for plasmid cloning in E. coli or yeast mating. We first validate MP3-seq using well-characterized coiled-coil heterodimer interactions and synthetic binders for different human BCL2 protein family members. Then, we apply MP3-seq to characterize interactions between rationally designed 2×2 and lx2 heterodimers and demonstrate that MP3-seq can measure over 100,000 PPIs in a single experiment. We identify successful designs and use a greedy algorithm to find potentially orthogonal subsets. We delve into the elements expected to confer 2×2 interaction specificity by screening variations of a successful 2×2 pair with MP3-seq. Finally, we predict complexes for coiled-coil dimers with monomeric AlphaFold2, and versions 1 through 3 of AlphaFold-Multimer. We assess predicted error values (iPTM, pLDDT, pAE) ability to predict coiled-coil interactions and orthogonality, and use said error values and physics-based structure energy terms from Rosetta to train models to predict MP3-seq values and classify interactions.

## Results

### MP3-seq workflow

In MP3-seq, all molecular components required for measuring interactions between two specific proteins are encoded on a single plasmid **(Figure 1A)**. A plasmid library is constructed directly through homologous recombination in yeast to measure all possible interactions for a set of proteins. To do so, we transform haploid MATa-type yeast with a mixture of DNA fragments. One of these fragments is a backbone carrying a centromere sequence, the selection marker, a DBD, and an AD. On average, the centromere sequence ensures that only one pair of proteins is expressed in each cell (Scanlon et al., 2009). We use a Cys(2)His(2) zinc-finger domain of the mouse transcription factor Zif268 (Mcisaac et al., 2013) as the 080 and its corresponding promoter to drive the expression of the growth essential enzyme *his3*. The herpes simplex virus-derived protein domain VP16 is used as the AD. Additional fragments contain one of the proteins of interest and its associated unique barcode separated by a terminator sequence. In the experiments described here, these fragments are ordered as a single oligonucleotide, but for longer proteins a barcode can be added to a protein of interest by PCR. Due to their short length and distinct sequences, we used Tsynth23 and Tsynth27 as terminators in most experiments (Curran et al., 2015).

**Figure 1.**
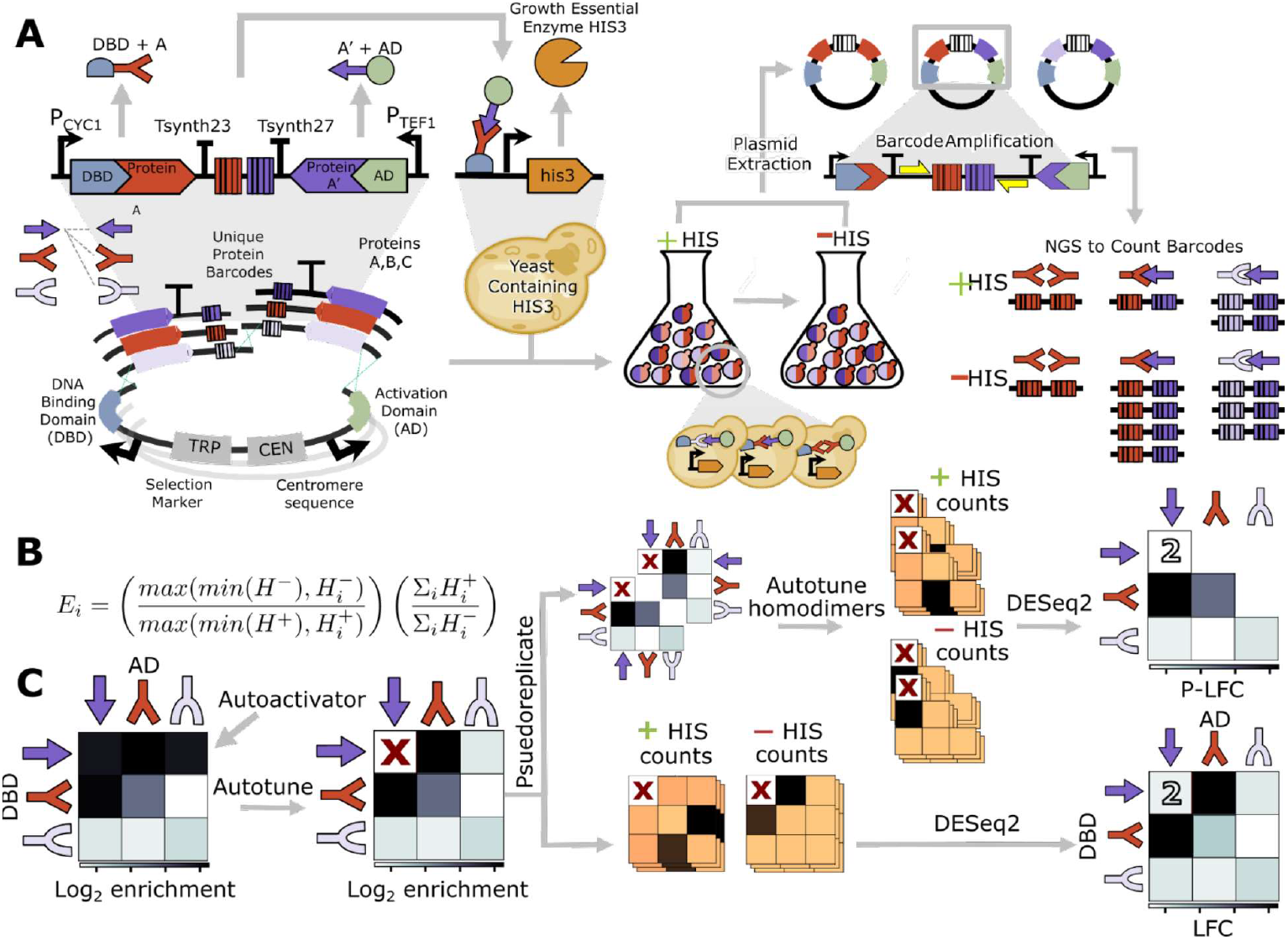
MP3-seq workflow. **A**. Barcoded DNA fragments encoding proteins to be screened, a fragment encoding two terminators, and a backbone fragment bearing a centromere signal, a DBD, and an AD are transformed into yeast cells via electroporation. Homology of fragment 5’ and 3’ ends allows plasmid assembly in yeast. The centromere sequence ensures the expression of one pair of hybrid proteins per cell on average. The cells are split into two populations and one is subjected to growth selection in media lacking histidine. If the hybrid proteins interact, the DBD and AD form a transcription factor that drives HIS3 expression. Therefore, interacting pairs should be enriched in the population post-selection. Cells are lysed, and barcode-containing fragments are amplified and sequenced. **B**. Enrichment can be calculated for each PPI i using library size normalized read counts with a pseudo count of the minimum detected value per condition. **C**. After enrichment calculation, each replicate is screened for autoactivators and corrected with Autotune. Replicate pre- and post-selection barcode counts are merged directly with DESeq2 or split into pseudoreplicates and then merged.

After transformation, we select in media without Tryptophan (Trp) to ensure plasmid maintenance. An aliquot of the cells is frozen, while the rest undergo selection in media lacking histidine (His). Plasmid DNA is extracted from pre-and post-His selection cells and the barcode-containing regions are amplified **(Figure 1A**, right). The barcode-barcode amplicons are sequenced, and their relative enrichment can be calculated from barcode counts to serve as a proxy for interaction strength **(Figure 1B)**. Some proteins’ expression or folding may be impacted by fusion with either the AD or DBD; therefore, we test all proteins fused to both the AD and DBD. Therefore, each PPI screened has two fusion orders (DBD fused to Pl, AD fused to P2 or DBD fused to P2, AD fused to Pl).

### MP3-Seq analysis pipeline

First, we calculate the enrichment between the pre and post-His selection stages for PPI fusion order. **(Figure 1B)**. Next, we detect autoactivators and replace their post-His selection values with values calculated from the non-autoactivating fusion order (see Autotune in Methods); autoactivation is an error mode in Y2H experiments where high enrichment is observed for a protein for all interaction partners, suggesting non-specific activation of selection marker expression (Shivhare et al., 2021). Typically, this behavior is observed fused to either the AD or the DBD but not in both orientations. Note that if an autoactivator is found its homodimer is not recoverable with Autotune as there is no reverse fusion order His measurement to use for correction.

Following autoactivator removal, enrichment values can be averaged together to obtain interaction strength values. Alternatively, to correct for variation in the read count distribution between experimental replicates resulting from different sequencing depths, selection times, and other experimental factors, we calculate log_2_ fold change (LFC) using the DESeq2 package (Love et al., 2014). DESeq2 calculates differential enrichment across multiple replicates and provides Hochberg-adjusted Wald test p-values, identifying PPIs with LFCs significantly different from an LFC of zero.

In some cases, it is desirable to combine MP3-seq measurements for both fusion orderings (e.g., Pl-DBD + P2-AD and P2-DBD +Pl-AD-> Pl-P2). For example, if comparing MP3-seq LFCs to Kds collected for the pair of interest (Pl-P2). The simplest method for merging these measurements is applying an operation to both, such as averaging or taking the max of the values. Here, we use an alternative approach and treat each fusion order as an independent set of measurements, turning them into two pseudoreplicates. These pseudoreplicates are combined with DESeq2 to calculate LFCs **(Figure 1B)**. We refer to these values as pseudoreplicate LFCs (P-LFCs).

### MP3-seq benchmarking with orthogonal coiled-coil dimers

To validate MP3-seq, we screened 144 pairwise interactions between six orthogonal 1×1s in the NICP set (Lebar et al., 2020) **(Figure 2A)**. A pool of 24 DNA fragments where each of the 12 proteins was fused to either the AD or DBD was ordered. Then, all interaction pairs were assembled in a pooled experiment, and interaction strengths were quantified with MP3-seq. We confirmed that selections were occurring as expected by checking pre-His selection barcode counts remained steady over time **(Figure S1A)**, and that applying different levels of 3-Amino-1,2,4 triazole (3-AT), a competitive inhibitor of His3, affected post-His selection counts **(Figure S1B-D)**. LFCs for all orientations were calculated from three experimental replicates, with interactions occurring almost exclusively between intended (on-target) partners (i.e., Pl:P2, P3:P4, etc., **Figure 2B)**. Homodimer plasmid assembly is less efficient in MP3-Seq and generally results in lower input barcode counts for these PPIs, but coverage was sufficient for inclusion in the analysis **(Figure S1E)**. This phenomenon is likely due to increased sequence homology, which interferes with DNA fragment order during plasmid assembly. For a more quantitative validation, we correlated MP3-seq LFCs with luciferase expression assay interaction scores in HEK293T cells from (Lebar et al., 2020) and found good agreement (R^2^=0.83, **Figure 2C). Figure 2D** represents P-LFC MP3-Seq data for these PPIs as a graph, where each protein is a vertex and edges are significant measured interactions weighted by P-LFC. This graph demonstrates one of the advantages of having adjusted p-values (p_adj_) values available to filter data to only higher confidence measurements for understandability.

**Figure 2:**
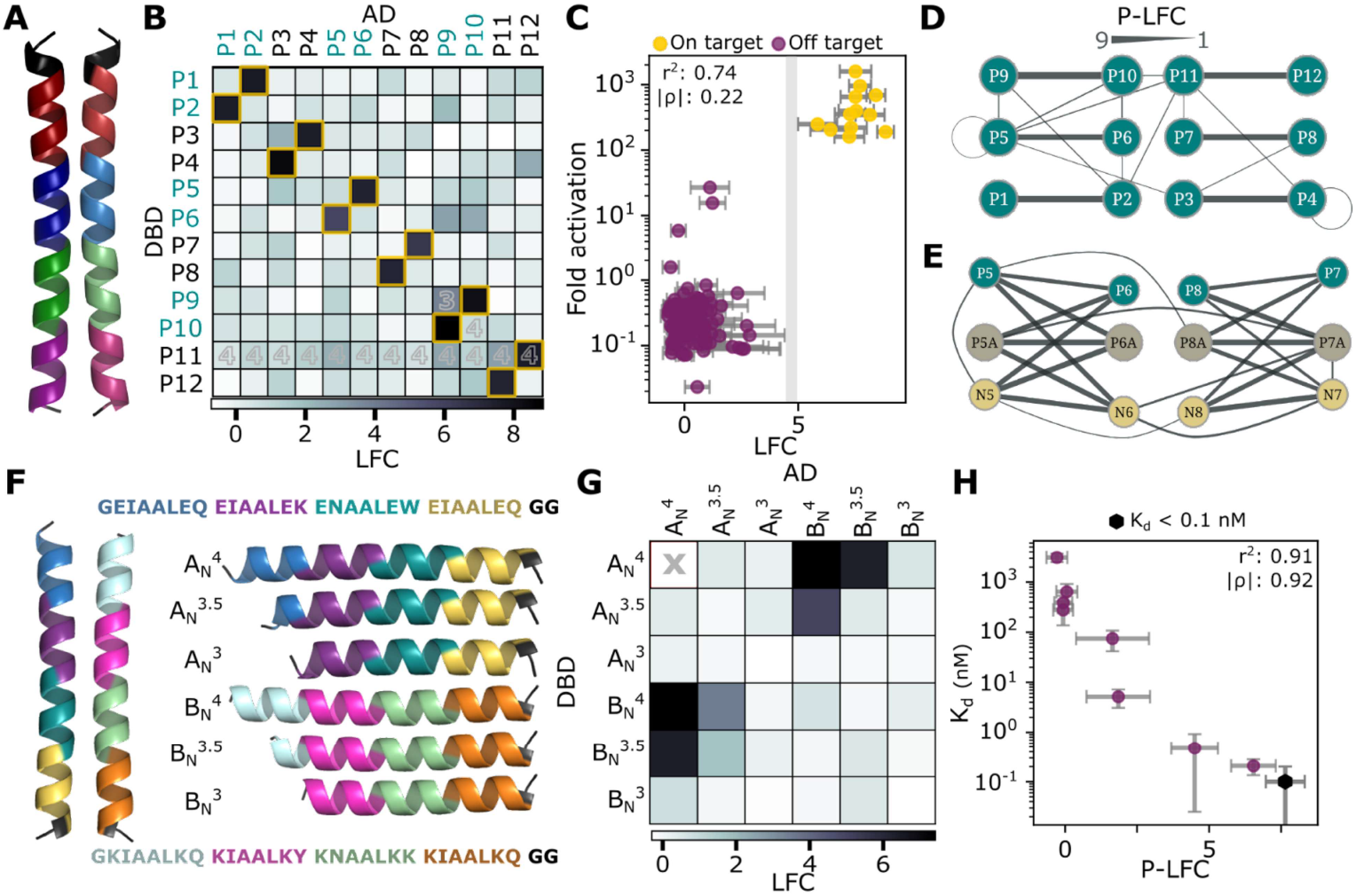
Validation of MP3-seq with coiled-coil heterodimers. **A**. NICP-series 1×1s from (Lebar et al., 2020) with complementary heptads designed to interact shown in like colors. B. MP3-Seq LFC (log_2_ fold change) of the NICP-series interactions. All values were calculated from 5 experimental replicates except for those labeled, where labels indicate the number of replicates available. Yellow outlines denote designed on-target interactions. **C**. Correlation of the on- and off-target NICP-series MP3-seq LFCs with fold activation fluorescence values from (Lebar et al., 2020) The gray bar is the gap separating on- and off-target interactions. Error bars are standard error. **D**. Filtering NICP-series interactions and **E**. P, PA, and **N** series designed coil interactions (Plaper et al., 2021) to include only those with P_ad_i 0.05. Line weights correspond to MP3-Seq P-LFCs. F. A designed 1×1 and its truncations from (Thomas et al., 2013) **G**. MP3-Seq enrichment for the A_N_ and B_N_ coils and their truncations with three replicates, gray ‘x’ indicates missing interaction. **H**. Correlation of MP3-Seq P-LFCs with K_d_ values from (Thomas et al., 2013) for the A_N_ and B_N_ truncations, horizontal error bars are standard error, vertical are standard deviation.

In a separate all-by-all experiment with 28 proteins, we screened the NICP 1×1s and two sets of 1×1s derived from them (N1, N2, N5-N8, and P5A-P8A) but with increased thermodynamic stability of their on-target interactions (Plaper et al., 2021). In this experiment, we expect related coils to have similar interaction patterns (e.g., PSA can bind to P6A, N6, and P6), which agrees with the interaction graph created from significant MP3-Seq P-LFCs in **Fig 2E**. The full interaction graph can be found in **Fig S2A**, where we show minimal off-target cross-talk between NCIP proteins and their variants. We exhibit low correlation with melting temperatures collected for the interactions by (Plaper et al., 2021), but good correlations with split-luciferase and split transcription factor interaction assays for this validation set **(Fig S2B-D)**.

To assess if MP3-seq values provide quantitative information about interaction strength, we measured interactions between a set of varying-length 1×1s designed to span a wide range of K_d_ values (Thomas et al., 2013) **(Figure 2F)**. We performed three all-by-all replicates; these interactions are not expected to be orthogonal as (Thomas et al., 2013) achieved weaker interactions by truncating the two parent four-heptad binders (A_N_^4^, B_N_^4^) by a half or full heptad **(Figure 2G)**. The MP3-seq P-LFCs correlated very well with previously measured dissociation constants over approximately three decades (r^2^=0.94, **Figure 2H)**.

We explored using minimum readcount filtering at different thresholds, and found that filtering brings minimal or no changes to correlation with the (Thomas et al., 2013) K_d_ values. However, for the NICP series, it does improve correlation (particularly for Spearmans’ rho) **(Fig S3A-B)**. Finally, to examine the performance for weaker interactions, we compared the correlation of enrichment and P-LFC values with the (Thomas et al., 2013) K_d_ values. We found that enrichments and P-LFC values had high correlations with all data points and similar intermediate correlations when only considering the five weakest interactions **(Fig S3C)**.

### MP3-seq benchmarking with BCL2 family binders

To validate MP3-seq outside of 1×1 interactions, we tested a set of proteins previously characterized by biolayer interferometry (Berger et al., 2016) and Alpha-seq (Younger et al., 2017) composed of six homologous proteins from the BCL2 family (Bcl-2, Bcl-xL, Bcl-w, Mcl-1, Bfl-1, Bcl-B) and nine *de nova* designed inhibitors of said homologs. A crystal structure of one of the synthetic binders bound to its target is shown in **Figure 3A**, while **Figure 3B** shows the domain organization of the six human proteins. All inhibitors were designed to interact with the BH3 domain binding pocket of the homologs. To better compare with the HT-Y2H method Alpha-seq, a truncated version of Mcl-1 was used.

**Figure 3:**
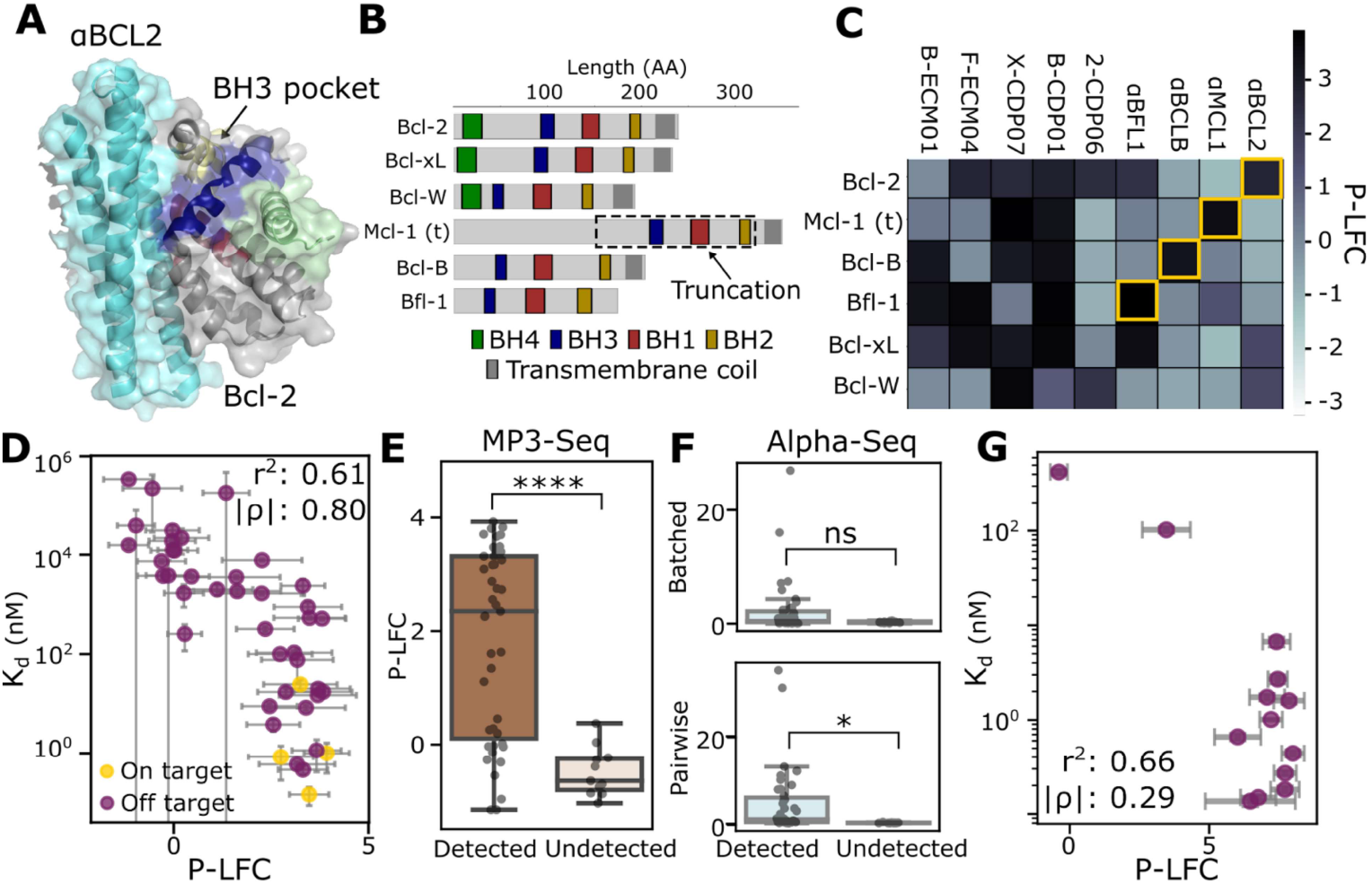
Validation of MP3-Seq with BCL2 proteins. **A)** Colored crystal structure of Bcl-2 and its designed BH3 binding inhibitor from (Berger et al., 2016) (PDB: SJSN). Binder in blue, Bcl-2 in gray with different domains in the binding region highlighted in color. **(B)** BH3 binding domain annotations of the six human Bel homologs measured by (Berger et al., 2016; Younger et al., 2017) with the experimental truncation used by Younger and in this work shown. **C)** MP3-Seq P-LFCs for inhibitor and Bcl-2 interactions. Yellow boxes highlight intended on-target interactions **D)** Correlations with K_d_ measurements from biolayer interferometry. **E)** MP3-Seq P-LFC **(F)** and Alpha-Seq distributions for interactions that were undetected by biolayer interferometry due to instrument detection limits (K_d_ 25,000 nM). **G)** Correlation of PUMA peptide and Mcl-1 (t) PPIs with PUMA peptide and full Mcl-1 K_d_ measurements from (Rogers et al., 2014).

An MP3-seq screen of all BCL2 homologs against all inhibitors is shown in **Figure 3C**. Version 1 of MP3-seq was used for the first two replicates of this experiment. Unlike version 2, barcodes were inserted into the 3’UTRs of the binders upstream of the terminators and could thus differentially impact mRNA stability and protein levels. However, the inter-replicate correlations suggest consistent interactions between versions **(Figure S3A)**. Only binders beginning with alpha correspond to the final, specific designs of (Berger et al., 2016), while the others are intermediate or failed designs. This can be seen in **Figure 3C**, with a divide between the largely orthogonal rightmost four columns of the heatmap of MP3-seq P-LFCs and the left, less specific columns.

Our data agree well with dissociation constant measurements obtained from biolayer interferometry by (Berger et al., 2016) (r^2^=0.61, **Figure 3D)**. We exhibit good agreement with Alpha-Seq percent survival for their low-throughput pairwise assay and high-throughput batched assay on the same interactions (batched r^2^: 0.45, paired r^2^: 0.61, **Figure S3B-E)**. Different measurement conditions can partially explain variation between our results and those published earlier. MP3-seq interactions are measured with proteins expressed in yeast, while biolayer interferometry uses purified proteins, and Alpha-seq displays proteins on the yeast surface.

Some BCL2 inhibitors failed to produce K_d_ values when measured with biolayer interferometry, likely because the interactions were weak and below detection limits. We examined the undetected (n=11) versus detected interactions (n=43) and found that the mean MP3-seq value of PPIs that biolayer interferometry detected was significantly greater than those which were undetected (independent t-test, Ha: μ_detected_ > μ_undetected_) **(Figure 3E)**. Pairwise Alpha-Seq also had the mean of detected interactions significantly greater than undetected ones. However, batched Alpha-Seq, the high throughput version of the assay, did not exhibit this behavior **(Figure 3F)**. Additionally, MP3-Seq correlation improves for batched but not paired Alpha-Seq when including all data points (batched r^2^: 0.45, paired r^2^: 0.69) **(Fig S3B-E)**. Together, these results show that MP3-seq can work with globular proteins in addition to coils and that MP3-seq results agree with those obtained by Alpha-seq and biolayer interferometry. The significant difference between the detected and undetected interactions provides evidence that MP3-seq excels at separating very weak PPIs from others at high throughput.

Finally, we screened a set of PUMA peptides to validate MP3-Seq performance for peptide-protein and disorder-dependent interactions. PUMA is an intrinsically disordered protein that binds to Mcl-1 at the BH3 pocket **(Figure 3B)**, where Mcl-1 binding results in the disordered PUMA BH3 binding motif transitioning to an ordered helix. Peptides of BH3 motif mutants were used to investigate the effects of helicity degree on this IDP-dependent interaction (Rogers et al., 2014). As part of this investigation, stopped-flow fluorescence techniques were used to measure the Kd values of a set of PUMA peptides interacting with the full Mcl-1 protein. We screened a subset of these peptides against our Mcl-1(t) protein and found a good level of correlation (r^2^: 0.66) **(Figure 3G)**.

### Large-scale assay of designed heterodimers

To explore the feasibility of using MP3-seq for large-scale screening, we prepared a library of designed heterodimers (DHDs) following (Chen et al., 2019). On-target pairs were designed as three- or four-helix bundles with buried hydrogen bond networks (HB-Nets) connecting all helices and then split into a helix-turn-helix hairpin and a single helix (1×2s) or 2×2s. Initially, we chose 100 designs for testing (DHD1, **Figure 4A)**, later adding 52 more designs in a second round of experiments (DHD2, **Figure 4A)**. As controls, we selected nine previously tested 2×2s (Chen et al., 2019). We also included 35 pairs of binders derived from these controls by modifying the hairpin loops or truncating the helices (DHD0, **Figure 4A)**. Finally, we added a series derived from a common parent 2×2 through truncating heptads (mALb, **Figure 4A)**. For each on-target pair, one protomer is designated “A” and the other “B”. Interactions between and within these groups were tested in MP3-seq experiments of varying sizes. We performed two replicates of an all-by-all screen including DHD0, DHD1, DHD2, and mALb proteins as well as the BCL2 homologs and cognate binders described above, resulting in a matrix of 337×337 =113,569 interactions. We also performed three replicates of a screen using only DHD1 with a subset of DHD0 and one with DHD0, DHD2, and mALB. MP3-seq version 1 was used for most experiments in this section. Still, the experiment highlights the large experimental scales achievable with MP3-seq and the inter-replicate correlations suggest that the data is internally consistent **(Figure S2 B-C)**.

**Figure 4.**
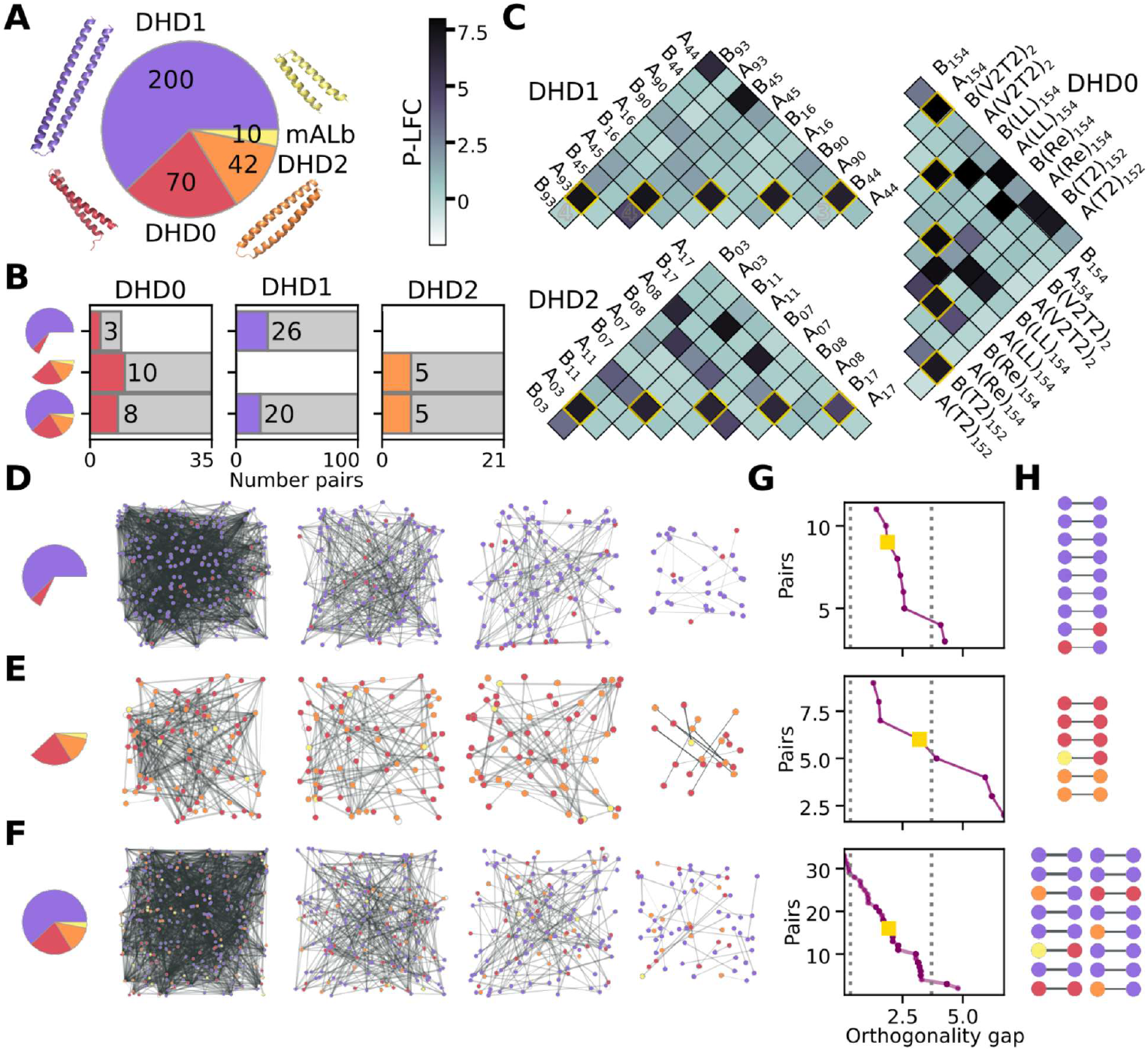
High throughput screening of designed heterodimers. **A**. Summary of the synthetic binder types. **B**. Successful designs for the three main sets. Gray bars are possible pairs for the set, and colored bars indicate the number of successes. **C**. Top 5 successful designs from each set by P-LFC. Yellow boxes show on-target interactions. **D**. From left to right, pre-DUET positive, p_adi_ 0.01 P-LFC PPI network for the DHD1 and DHD0 subset, and the network at DUET iterations 50, 75, and 113. E. Pre-DUET network for the DHD0, DHD2, and mALb sets, and the network at DUET iterations 10, 15, and 37 (final). F. Pre-DUET network for all designs and the network at DUET iterations 50, 75, and 114. **G**. Orthogonality gaps of the DUET final networks without significance filtering. Left and right dashed lines show the MP3-seq orthogonality gaps for BCL-2 and final inhibitors and the NCIP-series, respectively. Yellow squares correspond to half of the starting networks remaining. **H**. DUET final networks reduced to half their final iteration size.

We applied autotune corrections and symmetrized the read counts to calculate P-LFCs for all possible pairs. On-target interactions, where the A and B protomers were designed to interact, were evaluated for success. In this case, we defined a ‘success’ as any PPI with a p_adj_ ≤ 0.01 and a P-LFC ≥ 4. The number of successes out of the total possible on-target PPIs in each design set can be seen in **Figure 4B**. Every design set had an approximately 20% success rate in the largest screen, consistent with past 22% success rates for alpha-helical bundles (Ljubetič et al., 2017a). The top 5 P-LFC on-target interactions for each design category are shown in **Figure 4C**. The DHD pairs have strong on-target interactions. However dimers are not necessarily limited to the designed on-target pairs, even when examining only the PPIs of the A and B protomers of the top five designs.

### Finding orthogonal subsets

For all four sets of designs (DHD1-3, mALB), we detected a large fraction of strong off-target interactions (e.g., Al and B2 of DHD1 instead of A1 and B1). For example, in the all-by-all screen with all of DHD1-3 and mALB, only 33 of 914 total PPIs with p_adj_ ≤ 0.01 and P-LFC ≥ 4 were on-target.

Given our many measured strong interactions, we asked if we could extract potential orthogonal PPI subsets from our data. To this end, we used significant (alpha = 0.01), positive P-LFC values to construct a weighted undirected graph, as in **Figure 2D-E**. This approach allows us to rephrase the problem as finding a graph of degree one vertices without self-edges which maximizes the sum of the remaining edges. To do this, we developed a simple scoring function that rewards graphs based on existing orthogonal edges or those over a desired orthogonality gap and punishes graphs for non-orthogonal edges. This scoring function was used in a greedy graph reduction method, Deleting Undirected Edges Thoughtfully (DUET), which removes a vertex and its associated edges each iteration until a 1-regular graph remains (see Methods). We applied DUET to our different data sets (the number of degree one vertices and score of total graphs per algorithm step can be seen in **Fig S5A-B)**. For DHD1 and the DHD0 subset, we went from 2,001 edges between 202 vertices to 18 DUET pairs **(Figure 4D)**, for the DHD0/DHD2/mALb data from 279 edges between 85 vertices to 11 DUET pairs **(Figure 4E)**, and for all designs (DHD0-2,mALb) from 1,562 edges between 270 vertices to 36 DUET pairs **(Figure 4F)**. Of these, 2, 2, and 4 DUET pairs were on-target for DHD0/DHD1, DHD0/DHD2/mALb, and all designs, respectively.

The DUET final results are only orthogonal if all non-highly significant interactions are considered non-interacting. As this may not be the case, we used all P-LFC >0 between DUET pair protomers regardless of significance for a more conservative analysis. First, we removed interactions with protomers for which the DUET pair P-LFC was lower than the highest non-DUET pair P-LFC. Then, we reduced the remaining DUET pairs by removing whichever pair had the largest non-DUET P-LFC one by one **(Figure 4F)**. When reduced to half the starting DUET pairs (squares in **Figure 4F)**, the orthogonal sets shown in **Figure 4G** are left, which all have orthogonality gaps comparable to the NICP set in **Figure 2** and the BCL2-inhibitor pairs in **Figure 3**. An additional undesirable behavior for these potentially orthogonal sets would be if DUET is biased toward selected proteins that have missing interaction data between one another (and, therefore, fewer edges in the graph). To determine if this was the case, we ran permutation tests where we counted the number of missing interactions between the proteins in the initial DUET results and randomly sampled protein sets of the same size. We did not find that DUET results had a significantly higher number of missing interactions **(Fig S5C)**.

### Elucidating the rules of specific helix-loop-helix binding

High-throughput, high-quality data can be used to probe novel design rules to improve protein design. We demonstrate how MP3-Seq can contribute towards a better understanding of binding specificity in designed 2×2 pairs by testing variants of different lengths and with a different number of buried hydrogen bond networks. Our target pair, mALb8, is a high-affinity 2×2 heterodimer with three HB-Nets between its A and B protomers **(Figure SA)**. This pair was the basis of the mALb set introduced earlier.

First, two mALb protomer truncations were designed, removing one or two turns from the end of each alpha helix **(Figure 5B)**. Truncation 1 (T1) versions are one turn shorter than the original monomers, while Truncation 2 (T2) versions are two full turns shorter **(Figure S6A)**. The original binders and the truncations were screened in an all-by-all screen with MP3-Seq and LFCs were calculated from four replicates. It can be seen from **Figure 5B** that truncating the binders by two turns (T2) eliminates binding. The relationship between length and binding affinity is highly non-linear, as binding disappears once the minimum length is reached. We can use this data to infer the shortest possible length (3.5 heptades) for 2×2 binders containing two buried HB-Nets. This pattern is similar to that seen for the A_N_ and B_N_ single helix truncation experiments in **Figure 1D-E**.

**Figure 5:**
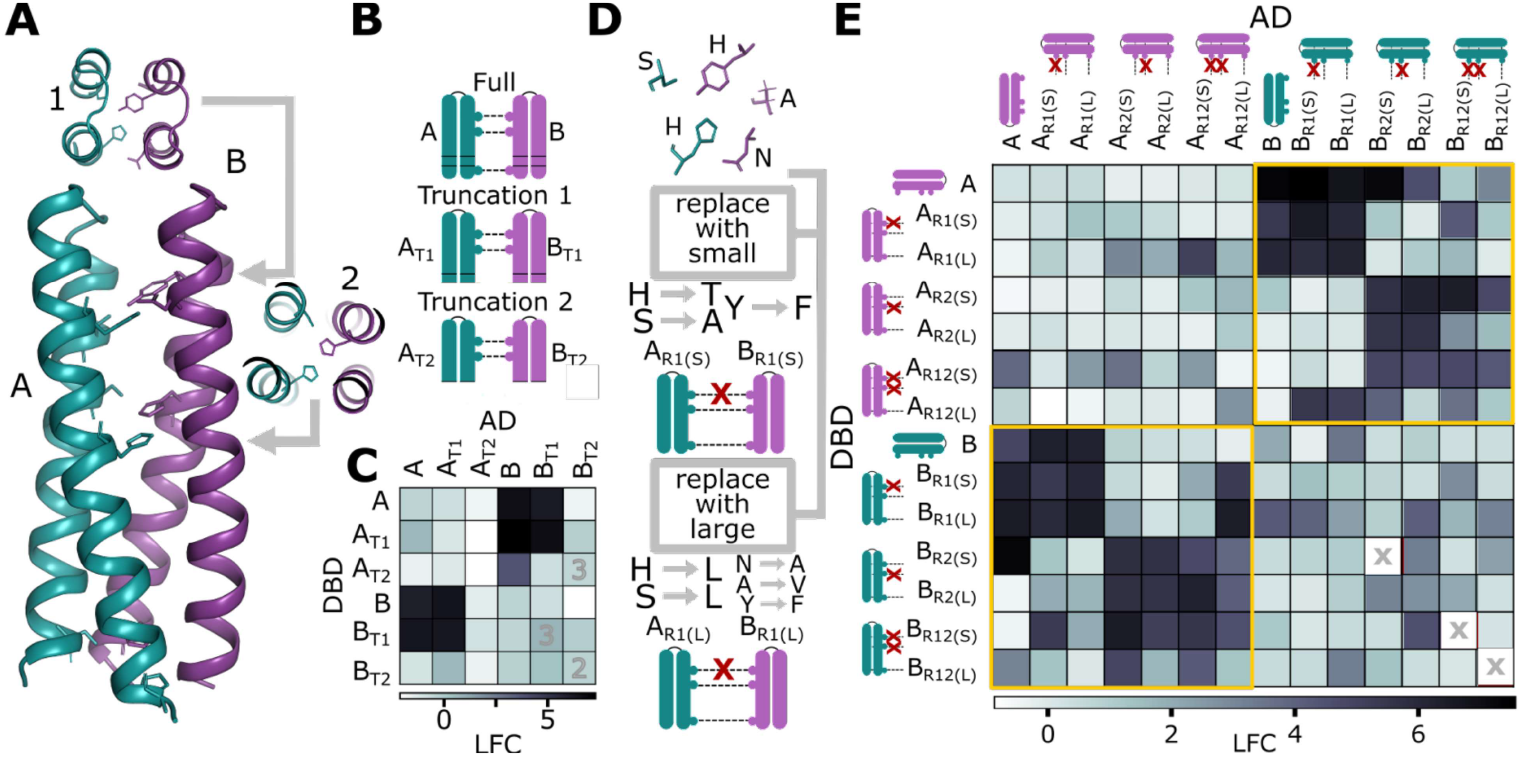
Systematic exploration of length and hydrogen bond network presence on helix-loop-helix binding. **A**. The mALb8 helix-loop-helix heterodimer with HB-Nets 1 and 2 shown. B. mALb8 and the two shorter variants (T1; one helical turn shorter, T2; two helical turns shorter). C. Log fold change (LFC) of the truncated mALb8 interactions. Gray X squares indicate insufficient reads. We observed a markedly non-linear response to the length of the binder. T2 variants do not bind. All PPIs had four replicates, except those marked with their replicate number. **D**. An example of HB-Net removal using the small and large hydrophobic replacement protocols for the first HB-Net E. LFCs of mALb8 with and without removed HB-Nets. S and L designate the replacement protocol used, while R_11_ R_2_, and R_12_ denote if the first, second, or both HBNets were replaced. Gray X indicates insufficient reads. The yellow boxed regions show that two HB-Net mismatches are enough to specify orthogonality. Hydrophobic residues alone cannot confer specificity.

To probe the specificity conferred by the buried HB-Nets in the 2×2 binders designed with the approach from (Chen et al., 2019), we have removed either the first, the second, or both buried HB-Nets from the mALb8 dimer. Removal was done by replacing HB-Net amino acids in both A and B protomers with hydrophobic residues that can not form hydrogen bonds. Two Rosetta-design replacement protocols were used: one using smaller (S) and one larger (L) hydrophobic residues for replacement. The networks are shown in **Figure SD**, while all replaced designs can be found in **Figure S6B**. We observe that the designs largely do not homodimerize, as interactions between various A and B protomers are low. These results show that at least two hydrogen bond network mismatches are needed to confer specificity and orthogonality **(Figure 5E**, boxed areas). For example, A_R1_ and B_R2_ binding would bury half of the second HB-Net on protomer A and half of the first HB-Net on protomer B, while A_R12_ and B_R2_ binding buries only the first HB-Net on chain B. The binding data suggest that pairs with one HB-Net mismatch can still exhibit partial binding affinity: for example, B_R12(S)_ weakly binds to both A_R1_ and A_R2_. Additionally, hydrophobic residue size alone is not sufficient to confer orthogonality. For example, both B_R1(S)_ and B_R1(L)_bind to both A_R1(L)_ and A_R1(S)_, regardless of the size of hydrophobic amino acid residues. The all-by-all screen of original, truncated, and HB-Net removed mALb proteins can be found in **Fig S6C**, which shows that the first truncation mALb proteins have similar binding patterns as the originals, but that the second continues to have eliminated binding.

### Predicting orthogonal coiled-coil complexes and training ridge regressors and classifiers

To assess AlphaFold’s ability to predict interactions, we predicted complexes for all 6 1×1 NICP pairs displayed in **Fig 2B**. We used AF-M v1-3 and the monomeric version of AlphaFold2 (with chains connected by a virtual linker (Mirdita et al., 2022)). Five AF2 models were run per dimer, using both orders of feeding in input sequences. We compared error metric (plDDT, PAE, etc.) correlation with MP3-Seq LFC values and on and off-target classification for the complex predictors. The monomeric AF2 and v1 AF-M performed poorer than later AF-M versions **(Fig S7A-B)**. The interface Predicted TM score (iPTM) score averaged across our five predicted complexes was among the best-performing metrics. The AF-M v3 averaged across all 5 AF models for the NICP interactions can be seen in **Fig 6A**. The increase in performance for both correlation with LFC values and on/off target classification can be seen in **Figure 6B**, where v3 improves over v2 in both cases. However, when comparing the performance of the structure predictor error metrics with the 1×1 binding predictor iCipa (Boldridge et al., 2020), AF-M still is underperforming (particularly in the classification task). Other 1×1 binding predictor performances can be seen in **Fig S7D**, which similarly outperform AF-M metrics.

**Figure 6:**
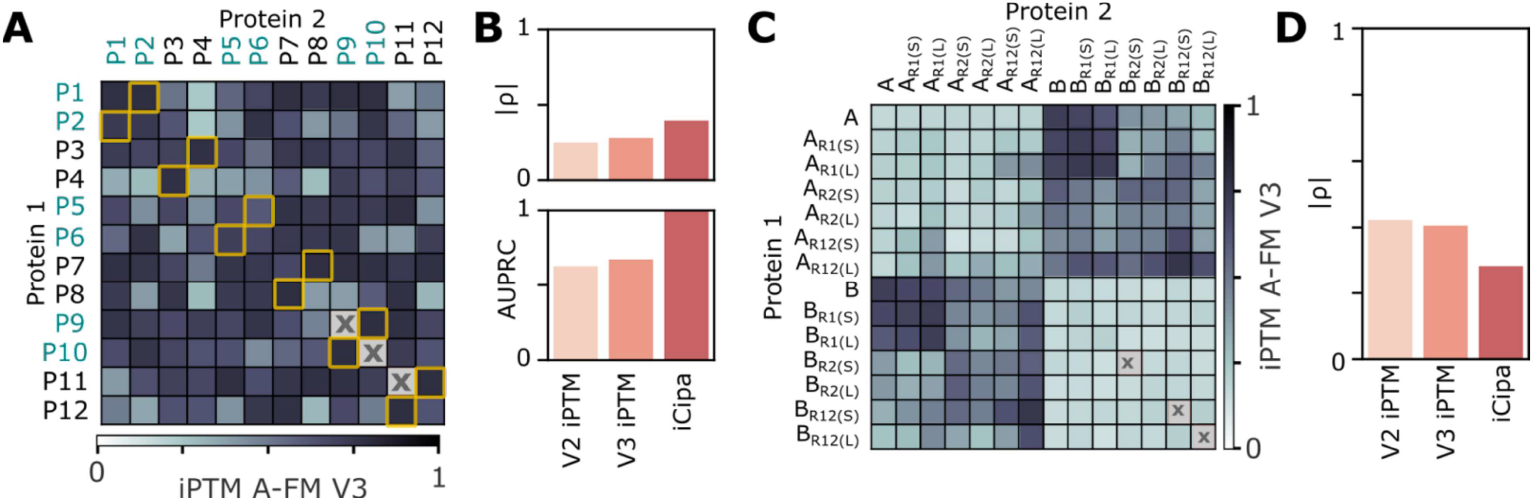
Predicted complex error for coiled dimers. **A**. Average predicted interface TM (iPTM) per complex from Alphafold2 multimer v3 (AF-M v3) for the **NICP** 1×1s. On-target interactions are marked with yellow squares. **B**. NICP correlation of average iPTM per complex with experimental **(Fig 2A)** LFC values (top) and area under precision recall curve (AUPRC) of classifying on- and off--target interactions (bottom). The 1×1 binding predictor iCipa (Boldridge et al., 2020) is shown for comparison. C. Average iPTM per complex with AF-M v3 for a subset of the mALb8 interactions. **D**. Correlation of average iPTM with mALb8 interaction LFCs **(Fig 2E)**. The 1×1 predictor iCipa is shown for comparison, and is unable to generalize to 2×2 coils unlike AF-M.

We then used the best-performing complex prediction models, AF-M v2 and v3, to predict structures for the truncated, mutated, and original 2×2 mALb8 complexes examined in **Fig 5**. There is no apparent increase in error metric LFC correlation with AF version numbers for this set as in the NICP set **(Fig S7F-G)**. The AF-M v3 averaged across all 5 AF2 models for a subset of mALb8 interactions can be seen in **Fig 6C**. Heterodimeric interactions have higher confidence than homodimeric complexes, but the effect of HB-Networks on orthogonality is not captured in detail. Although AF-M v3 iPTM captures some binding rules investigated for the mALb8 dimer, the iPTM values it does not have a clear advantage over v2 as in the NICP series. The better generalization ability of structure predictors can be seen in **Fig 6D**, where both AF-M versions outperform the 1×1 model iCipa when applied to 2×2 interactions. Therefore, while AF-M metrics may not outperform specialized binding predictors on the domain they were trained on, they have undoubtable cross-domain generalization abilities.

Encouraged by the demonstrated generalization abilities of AF-M, we set out to determine if the predicted structures can be used to predict LFC values and classify strong interactions. Rosetta was used to collect physics-based metrics (energy of interaction, surface of the interface, etc.) for each simulated dimer complex. Agglomerative hierarchical clustering was used to reduce the number of multicollinear features between the collected energy terms and AF error metrics **(Figure S8)**. Linear least square ridge regression models and logistic ridge regression classifiers were trained with on feature sets of decreasing sizes (see Methods).

We used two train-test approaches to investigate ways these models might be applied. For the first, we ranked all data points by their padj values and used them to partition the dataset into high-quality test set interactions and a mix of high and low-quality training set interactions. This approach was chosen to assess the ability of models using AF-M complex features to fill in missing interactions in an assay. An example test set can be seen in **Fig 7A**.

**Figure 7.**
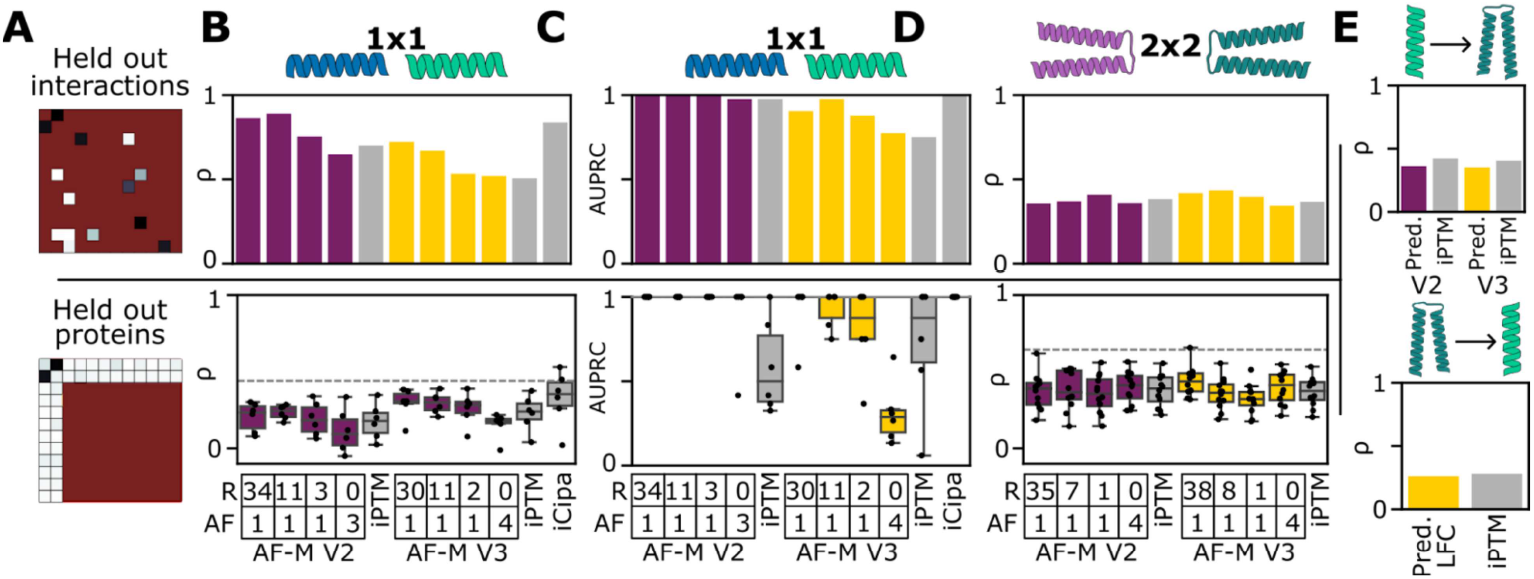
Training simple models on predicted dimer features to predict MP3-Seq values. **A**. Examples of the two different train-test splitting techniques: held out interactions, where half of the low MP3-seq P_ad_i LFCs are used for the test set (top) and held out proteins, where all interactions involving selected proteins are part of the test set (bottom). The dashed line shows the performance of the best held-out interaction model with the same number of training points as the held out protein models. Red interactions are train/validation, and non-red are test. Held out interaction (top) and held out protein (bottom) results, with features used for model training split into the number of Rosetta (R) and AF-M metrics (AF) used, for **(B)** predicting the NICP LFC values, **(C)** classifying the NICP interactions as on- or off-target, and **(D)** The mALb8 original, truncated, and HB-Net removed interaction LFCs. The dashed lines indicate the performance of the best held out interaction model with similarly-sized train/test set splits. **E**. (Top) Performance of predicting the LFC values of mALb interactions using the AF-M models with only AF inputs and comparing them with iPTM. (Bottom) Predicting the NICP LFC values with the AF-M v3 LFC prediction model.

In the other approach, proteins were selected, and all interactions involving those proteins were assigned to the test set. The protein-based split was used to evaluate how AF-M complex-based models could predict interactions involving new proteins instead of filling in missing interactions between known proteins (see Methods for how held-out proteins were selected). Multiple train/test sets were created for this split by holding out different protein sets to get a distribution of test performances. An example test set can be seen in **Fig 7A**.

The NCIP regression models performed similarly to iCipa for the held out interaction task and had gains over using iPTM alone for prediction **(Fig 7B**, top). In particular, the V2 and V3 AF-M models performed well, while the monomeric AF and AF-M V1 models were less successful, particularly when smaller training sets were used **(Fig S9A)**. A similar trend was seen for the held out interaction classification models, with the main difference being that the V2 iPTM provided an excellent classifier for this test set, and AF-M V2 performed better than V3 in modeling **(Figure 7D**, top). The number of Rosetta (R) and Alpha Fold (AF) features each model used can be seen in the tables at the bottom of **Fig 7B-D**, and more features yielded better models (notably when decreasing training set sizes, see **Fig S9A-B)**. All regression models dropped drastically in performance in the held-out protein task. However, this performance decrease can be partially attributed to a smaller training set, where the dashed line shows the performance of the best similar training set size held out interaction model **(Figure 7B)**. An additional thing to note is that in some cases, iPTM outperformed models using only AF error metrics on the test-set due to overfitting on the training set **(Fig S10C)**. The classification task’s performance stayed relatively high between the two tasks even when the training set size was reduced **(Fig S10A-B)**.

In contrast, the more complicated mALb interactions did not show clear trends in model performance with AF-M versions **(Fig 7D)**. It had a much smaller gain in performance in the held-out interaction task between the AF-only models and those with R and AF features. However, interestingly when dropping the training set size, there were gains in model performance, likely due to the held out interaction test set initially consisting of only high LFC interactions **(Fig S11)**. We then used our best AF-feature-only held out interaction NICP models to predict LFC values for the mALb interactions and vice versa **(Fig 7E, Fig S9C-F, Fig S118)**. We found in both cases that the models performed worse than iPTM when applied to coiled-coil interactions different from those they were trained to model.

## Discussion

Here, we introduce MP3-seq, an easy-to-use HT-Y2H method that can measure pairwise PPIs in a single yeast strain. In MP3-seq, the identity of each protein is encoded in a DNA barcode, and the relative barcode-barcode pair abundance before and after a selection experiment serves as a proxy for interaction strength. We also developed a data analysis workflow based on DESeq2 that makes it easy to merge replicates, remove autoactivators, and identify statistically significant interactions. The MP3-seq workflow could be further generalized in future work by adding autoactivator screening or adjusting the selection scheme to eliminate autoactivators (Shivhare et al., 2021). MP3-seq could also be combined with protein stability and expression assays to confirm that proteins tested in interaction screens are folded correctly and that a negative interaction measurement is not due to reduced protein levels. In particular, it is easy to imagine integrating MP3-seq with other growth selection-based workflows such as Stable-seq (Kim et al., 2013) or high-throughput protease assays (Rocklin et al., 2017). We expect stability screening to be more important for human protein than for the synthetic binders tested here, which are generally designed to have very stable folds.

We validated MP3-seq using several sets of proteins for which interactions had been previously characterized. In particular, we used a family of human pro-apoptotic proteins BCL2 and their *de nova* designed inhibitors, a set of peptides binding to MCL-1, and three different sets of coiled-coil peptides. We found quantitative agreement between our results and those reported previously. We then applied our method to characterize interactions in a pool of *de nova* 2×2 heterodimers that contain buried hydrogen bond networks. We showed that our approach could scale to measuring more than 100,000 interactions in a single experiment. Our computational workflow enabled us to identify potential orthogonal subsets, at various orthogonality gaps, within larger all-by-all screens.

Moreover, by screening interactions between protomers with truncations and modified hydrogen bond networks, we probed design rules for 2×2 binders with two buried HB-nets. Firstly, we showed that the minimum length for strong binding is 3.5 heptads and that binding affinity drops sharply for shorter designs. Next, we showed that at least two hydrogen bond network mismatches are needed for orthogonality, thereby setting minimum requirements for future sets of orthogonal 2×2 binders with buried HB-nets.

Finally, we assessed the structural prediction models AlphaFold2 and AlphaFold-Multimer to predict coiled-coil interactions. On the set of 1×1 dimers, we found that a sequence-based predictor (iCipa) correlated better with experimental LFC values than any single AF metric. However, AF-M metrics performed better than iCipa for the 2×2 dimers. Next, we tested if we could use a combination of AF2 and Rosetta metrics to improve AF2 interaction predictions. We found that regression models could be trained to fill in missing interaction LFCs from an all-by-all screen for the 1×1 dimers and, to a lesser extent, the 2×2 dimers. However, regression models performed poorly when test sets were selected to mirror the task of predicting interactions of ‘new’ proteins not present in the PPI training sets. In contrast, classification models performed well on filling in test interactions selected from the all by all matrix and classifying PPIs as strong, on-target interactions versus weak. None of our models performed well when predicting PPIs for the other coiled-coil group (predicting 2×2 interactions with the 1×1 models and 1×1s with the 2×2 models). Our results suggest that AF-M-predicted complex energy values may be used with minimal data to differentiate strong from very weak interactions for simple proteins, with the potential to expand to predicting interactions for new proteins unseen in the assay. Therefore, AF-M and AF-M complex prediction-based models could be valuable tools for qualitatively pre-screening interactions, but it remains necessary to measure interactions experimentally to determine their relative quantitative strengths.

We believe that the increased scale and streamlined workflow of MP3-seq will further accelerate the adoption of HT-Y2H methods and facilitate their use in additional applications. For example, high-throughput data collected with MP3-seq could be used to train predictive models of PPIs. Such models could be combined with generative algorithms to iteratively design heterodimers with desirable properties or with neural network interpretation to uncover determinants of binding specificity. Moreover, MP3-seq could be applied to characterize human PPIs or to screen variants’ impact on PPIs at high throughput.

## Acknowledgments

This work was supported by NIH Award R01GM120379 and ONR Award N00014-16-1-3189 to G.S. A.L. would like to acknowledge funding of European Commission MSCA CC-LEGO 792305 and Slovenian research agency project CC-TRIGGER Jl-4406.

## Data and code availability

Barcode count data and scripts necessary to recreate the MP3-seq pipelines, analysis, and models can be found at https://github.com/Seeliglab/MP3-DUET-AUTOTUNE

## Supplementary materials

### Methods

#### Experimental workflow

##### Plasmids and fragments

All protein binder coding sequences were ordered as double-stranded gene fragments from Twist Biosciences. In the gene fragment, the binder’s sequence is preceded by the coding sequence for a peptide linker, either the poly-Glycine-Serine (G4S) linker or the Glycine-Glycine Poly-Serine (GGS4) linker for the DBD and AD fusion proteins, respectively. This sequence also serves as a primer handle for initial fragment amplification and a homology sequence during plasmid assembly in yeast. The binder’s sequence is followed by a stop codon and a synthetic terminator from (Curran et al., 2015). Tsynth23 and Tsynth27 are used to terminate the transcription of DBD and AD fusion proteins, respectively. The terminators are followed by 21nt PCR handles for barcode amplification and predetermined 20nt long DNA barcode sequences. Barcodes were followed by 22nt long insulation sequences. This insulation sequence serves as a homology sequence during plasmid assembly in yeast.

The plasmid vector contains 30 nt end homologies with the G4S peptide linker and GGS4 peptide linkers in the binder fragments for assembly in yeast. The vector also contains the AD (VP16) and the DBD (zinc finger DNA-binding domain of mouse transcription factor Zif268). An ColE1 origin and beta lactamase expression for antibiotic resistance are also included to enable cloning in E. coli for troubleshooting. The vector uses a CEN/ARS sequence for yeast replication, expressing the plasmid genes on a stable, independent mini chromosome. TRPl expression is included for tryptophan selection.

##### Competent Yeast Cell Preparation

The yeast transformation protocol from (Benatuil et al., 2010) was adapted for this workflow. *5. cerevisiae* cells (EBY100) were grown overnight to stationary phase (00600 around 3.0) in YPD media (10 g/L yeast nitrogen base, 20 g/L Peptone, and 20 g/L 0-(+)-Glucose) on a platform shaker at 225 rpm and 30 °C. The following morning, we inoculated 100 ml of YPD media with the overnight culture at an initiate 0.3 0D600. The inoculated cells were grown on a platform shaker at 30 °C and 225 rpm until 00600 was approximately 1.6 (or for about 5 hours). Yeast cells were collected by centrifugation at 3000 rpm for 3 minutes and washed twice with 50 ml ice-cold water and once with 50 ml of ice-cold electroporation buffer (1 M Sorbitol / 1 mM CaCl2). Cells were resuspended in 20 ml 0.1 M LiAc/10 mM OTT in a culture flask and incubated for 30 mins at 30 °C and 225 rpm. We conditioned the cells by collecting them with centrifugation, washing once with 50 ml ice-cold electroporation buffer, and resuspending in 100 to 200 μL electroporation buffer to reach a final volume of 1 ml. This corresponds to approximately 1.6 × 10^9^ cells/ml and is sufficient for two electroporation reactions of 400 μL each. The cell culture and preparation can be proportionally scaled up if more electrocompetent cells are needed to make larger or more libraries.

### Library Transformation

Prepared electrocompetent yeast cells were kept on ice until electroporation. We prepared the DNA by combining 4 μg of digested vector backbone and 12 μg of each DNA insert for each 400 μL electroporation reaction. The DNA mixture was reduced via precipitation and resuspension as necessary to reach a volume of less than 50 μL. We combined 400 μL of electrocompetent cells and the DNA mixture and transferred it to a pre-chilled BioRad GenePulser cuvette (0.2 cm electrode gap), then put it on ice for 5 minutes. Cells were electroporated at 2.5 kV and 25 μF. Typical time constant ranges ranged from 3.0 to 4.5 milliseconds. Electroporated cells were transferred from each cuvette into 8 ml of 1:1 mix of 1 M sorbitol: YPD media in a culture flask and incubated on a platform shaker at 225 rpm and 30°C for 1 hour. Cells were collected by centrifugation and resuspended in synthetic complete media lacking tryptophan amino acid, SC-TRP media, (20 g/L glucose, 6.7 g/L yeast nitrogen base without amino acids, 5.4 g/L Na_2_HPO_4_, 8.6 g/L NaH_2_PO_4_·H_2_O and 5 g/L casamino acids [CSM-TRP]). 250 ml SC-TRP media were used for every 400 μL transformation reaction. The optical density value OD600 of resuspended cells varied between 0.3 and 0.4 depending on the experiment. A small aliquot of the resulting cell suspension was diluted 10-^3^, 10-^4^, 10-s, and 10-^6^. We plated 100 μL of each dilution onto SC-TRP agar plates and incubated at 30 °C in a static incubator. Library size was determined from the colony counts on the agar plates after three days of incubation. The remaining transformed cells in SC-TRP media were grown in a baffled flask overnight at 225 rpm and 30 °C. The resulting transformed library was glycerol stocked the following day after about 20 hours of incubation at OD600 of 1.0 by combining 750μL of culture with 750μL of 50% v/v sterile glycerol in a cryovial. Glycerol stock was stored at -80°C.

### Library Selection

After the 20-hour incubation period above, the passage process was started by diluting 20-hour transformed library cell culture in a clean autoclaved baffled flask with fresh 250 ml of SC-TRP media at a 1:400 ratio and incubating at 30 °C and 225 rpm. Alternatively, the transformed library can be inoculated from a glycerol stock at the same dilution and final volume. The cells were grown to approximately 1.0 OD600 value at which point the dilution procedure was repeated as needed with the same parameters until 36 hours had passed.. This passage process allows yeast cells to drop redundant plasmid copies and achieve an average of a single plasmid per cell. At the end of the passage process, a fraction of cells was inoculated in a baffled flask with SC-HIS-TRP media (20 g/L glucose, 6.7 g/L yeast nitrogen base without amino acids, 5.4 g/L Na_2_HPO_4_, 8.6 g/L NaH_2_PO_4_·H_2_O and 5 g/L casamino acids [CSM-TRP-HIS]) at a 1:400 ratio to start selecting for interactions. The remaining SC-TRP cells were collected by centrifugation at 5000rpm (3000xg) for 5 minutes and resuspended in 200 μLof Zymo Research Solution 1 Digestion Buffer per 50 ml of cells and stored in -80°C freezer until plasmid extraction. At least 100-fold over library size number of cells were collected. The selection for interaction step usually lasted between 24 and 36 hours depending on the library and concluded when cells in the SC-HIS-TRP flask reached OD600 value of 1.0. At that point, cells were collected by centrifugation, resuspended in Solution 1 buffer, and frozen at -80°C.

### Yeast Plasmid Extraction Protocol

Cells after TRP selection and HIS selection were collected by centrifugation as described in the library selection protocol. Cells were resuspended in 200 μL Zymo Research Solution 1 Digestion Buffer per 50 ml of cell culture. Resuspended cells were frozen at -80°C. We then thawed the cells and added 20 μL of Zymolyase per each 200 μL volume of cells. Cells were then incubated at 37°C for 4 hours, mixing once per hour by inverting the tubes ∼ 20 times each. Cells were again frozen at -80°C and thawed. 200 μL of Zymo Research Solution 2 Lysis Buffer was added to each sample, inverted to mix, and incubated at room temperature for 3-5 minutes. We then added 400 μL Zymo Research Solution 3 Neutralizing Buffer to each sample and inverted to mix. Lysed cell were centrifuged at max speed (about 17,000 xg) for 5 minutes. Supernatant was transferred to a new microcentrifuge tube and spun again at 17,000 xg for 5 minutes. Supernatant was transferred to Zymo-Spin I Columns in a collection tube and centrifuged at 13000 xg for 30 seconds. Flowthrough was discarded. Columns were then washed by adding 550 μL of Zymo Research DNA Wash Buffer onto the column and centrifuged for 1-2 minutes at 13,000 xg. Flowthrough was discarded and wash was repeated. Empty column was spun for 1 min at 13,000 xg to dry. DNA was then eluted by adding 20 μL of molecular grade water to the column, allowing it to incubate for 5 minutes at room temperature, and spinning for 1 min at 13000 xg. Eluate was then reloaded onto the column and the spin was repeated.

#### Quantitative PCR (qPCR) and Next Generation Sequencing Preparation

Using plasmid as the template DNA, the first round of qPCR amplified the barcode region and added a 20nt long unique molecular identifier (UMI) and lllumina Read1 and Read2 primer sites. KAPA HiFi HotStart ReadyMix and suggested reagent and PCR cycle parameters were used. The annealing temperature was 67°C. The reaction was allowed to proceed until the qPCR curve reached an inflection point, and the resulting fragments were purified using KAPA Pure Beads. The resulting product was used as the template for the second stage of qPCR to add sample indexes and P5/P7 flow cell adaptor sites. This reaction was allowed to proceed for 6-7 cycles and used the same parameters as the previous qPCR reaction. An Agilent real-time PCR machine was used for all qPCR steps.

### Analysis Workflow

#### Barcode counting

Fastq files were obtained from lllumina NextSeq 550 or MiSeq sequencers using bcl2fastq with the default parameters. Cutadapt (Martin, 2011) was then used to trim out fastq barcode regions, and barcodes were clustered using Bartender (Zhao et al., 2018) with d=2.

#### Autoactivator screening

We defined autoactivators as proteins for which the trimmed interquartile mean of the library normalized enrichments is an extreme positive outlier (interquartile range> 3) with respect to the other assayed proteins. Autoactivator screening was carried out for both possible domain fusions.

#### Autotune autoactivator correction

Following autoactivator detection, the non-autoactivating alternate fusion ordering data was used to infill the pre-His selection read count according to Equation 1 and the post-His barcode read count according to Equation 2.

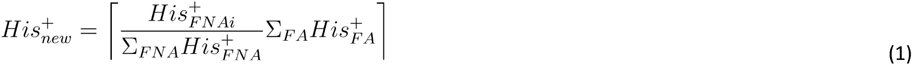

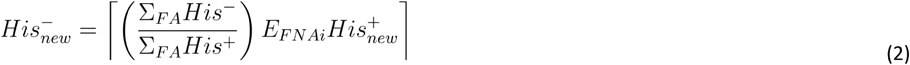

Where FA values are for the fusion order values which has autoactivation, FNA are for the fusion order values which does not have autoactivation, and *i* is the PPI in question. The summation of the His counts for selection are done pre-update of all new barcode count corrections. Note that since there is only set of values for homodimers there can be corrections made.

For Autotune of homodimers, Equation 3 is used to infill post-His selection values for both pseudoreplicates, using the same starting pre-His selection barcode count.

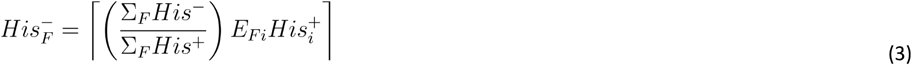

Where *F* is the fusion order being used to construct the pseudoreplicate, and i is the homodimer PPI. All autotuned proteins are available in **Table S1**.

### Undetected PPI filtering

For all experiments, if a PPI had a barcode count of 0 in the pre-His and post-His selection stages, it was considered undetected. Otherwise, if it was only undetected in one stage, the other was infilled using the corresponding minimum (missing pre-His reads were infilled using the minimum of detected pre-His reads, etc.). For any missing PPIs, the largest possible set of replicates were used during data processing for DESeq2 analysis.

### DeSeq2 Settings

DESeqDataSet objects were prepared with raw pre and post-His read counts, without infilling. A contrast using the pre and post-His columns as the conditions was used, and ashr lfcShrink was used to get final Mp3-seq LFC values. For pseudoreplicate MP3-seq, pre and post-His conditions were used along with a fusion order batch effect for a contrast of “‘cond + batch for DESeqDataSet creation. P-LFCs were obtained by ashr lfcShrink.

### DUET

A Pre-DUET weighted undirected graph *G* was created from MP3-seq values with P_adj_ ≤ 0.01 and P-LFC > 0, and vertices with degree 0 removed. Then, until I *G*I =0 or Δ(*G*)=l, all *G*\{v} were scored for *v* ∋ *V* with Equation 4, using $c=4$ and the highest scoring vertex removal was used to reduce the graph. The greedy reduction algorithm pseudocode can be seen in **Figure 6**.

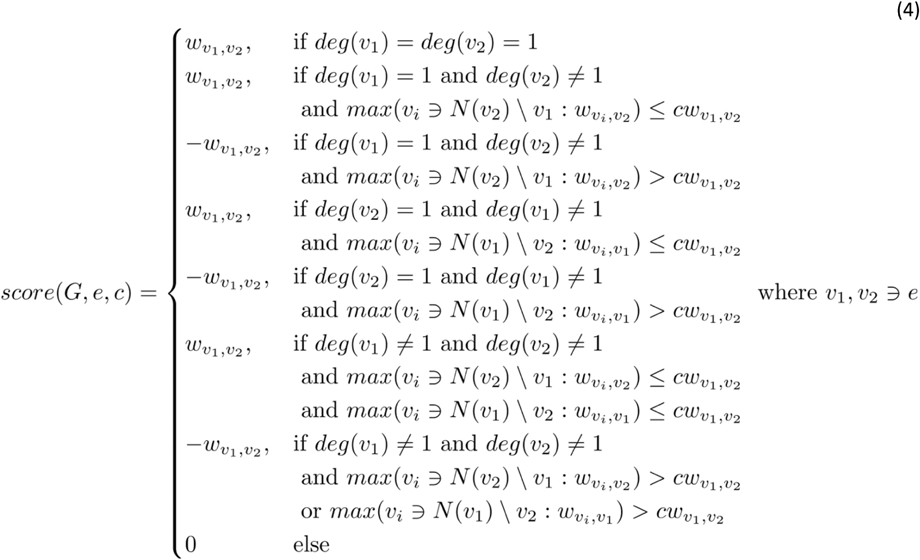

**Figure 6.**
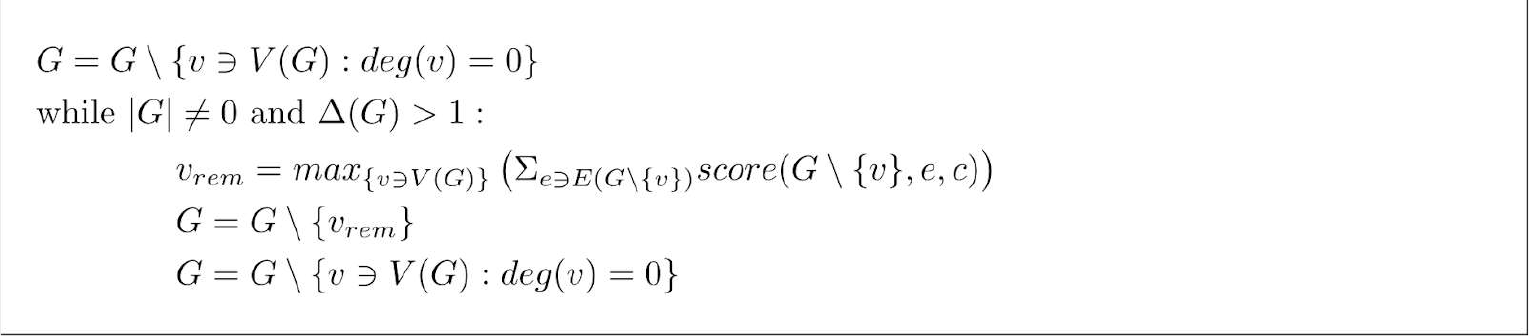
DUET pseudocode.

### Orthogonality gap reduction

To reduce the DUET pairs, any P-LFC>O interaction between the set of proteins involved in the DUET pairs was considered a real interaction. Non DUET-pair interactions were sorted by P-LFC, and the largest value interaction was selected to remove a DUET-pair. Whichever protein was involved with the smaller P-LFC DUET-pair and its DUET partner was removed. This was done one protein pair at a time until only one pair remained.

### Computational protein design

#### Design of helical bundles

Helical bundles were designed using methods developed in (Chen et al., 2019). Briefly, systematic sampling of Crick coiled-coil parameters was used to independently sample all helices of the heterodimers supercoiled around the same axis. The Rosetta HB-Net protocol (Boyken et al., 2016) was used to design buried HB-Nets in the core of the designs.

#### Hydrogen bond network removal

The design model of mALB8 was used as a starting point. The two hydrogen bond networks were manually annotated (R1 has residues 23+40+84+96, R2 has residues 16+43+103), see **Figure S5**. Two alternative protocols were used for each redesign. Either a FastRepack with a fixed static backbone, which results in replacement with smaller amino acid residues (T, A), or a FastDesign protocol, where the backbone was allowed to move, which resulted in replacement with larger amino acid residues (V, L, F). XML scripts for both protocols are included in the github repository.

The residues of the HB-net and all residues within a 6 A shell were redesigned, and a shell of 8 was repacked. The beta_nov2016 score function was used. The charge was fixed to -5 for each chain. Five sequences were made for each design. The sequences were filtered by score_per_res>-3.5, shape complementarity >0.7, sbuns<2, and ranked by ddg (the lowest ddg was chosen for the final design). Rosetta version 2021-07 (c04b624d89b45b55f573bbf2f6685ffbf2e4fcaf) was used for the calculations.

### AlphaFold structure prediction and predicted structure models

#### Structure Prediction

We used Alpha Fold 2.3.4 (Jumper et al., 2021) through the colabfold 1.5.2(Mirdita et al., 2022) on a local SLURM cluster. Alphafold was installed with the help of instructions from LocalColabFold (https://github.com/YoshitakaMo/localcolabfold). The following weights were used: alphafold2_ptm (6 recycles), alphafold2_multimer_v1 (6 recycles), alphafold2_multimer_v2 (6 recycles), alphafold2_multimer_v3 (auto stop recycles). Amber relaxation was used after the AF2 prediction. Five models were made per sequence pair.

#### Rosetta metric calculations

Rosetta version 3.13 (commit b3365a12916d9cl25d41) was used for the calculation of interface metrics. The energy function used was beta_nov16. Calculations were performed through the Rosetta Scripts XML (Fleishman et al., 2011). Two protocols were tested. In the first (min-sc) only the side chains were minimized. The MinMover with cartesian minimisation was run for 1000 cycles using the lbfgs_armijo_nonmonotone method with tolerance of 0.001. In the second (flex-bb) the backbone atoms were also allowed to move during minimisation. First side chains were minimized then the relative positions (jump 1) between the heterodimers and finally all atoms were minimized using the same parameters as above. Then the interface was selected using ResidueSelectors and the interface lnterfaceAnalyzerMover was used for the analysis. Additionally SASA (solvent accessible surface areas), ddG (difference of the energy between the bound and unbound state), shape complementarity, contact molecular surface and the number of buried hydrogen bonds (Coventry and Baker, 2021) were also calculated.

#### Collinearity feature reduction and feature engineering

To reduce the number of collinear features, all non-numeric features and features with only three or less unique values were removed from the set of AF and Rosetta metrics. The exception to this is the number of clashing residues, which was used to calculate the sum of clashing residues in the complex as a feature. Following this, Spearman’s correlation coefficient was calculated between all features, and used to make a distance matrix for hierarchical clustering using Ward’s linkage. A dendrogram of features was then created with these linkage values, and percentiles of the dendrogram’s total height were taken as thresholds to reduce the number of features until only two features remained. The results of this process can be seen in **Fig S8**. An additional ‘baseline’ model of only AF error metrics without collinearity reduction was also prepared. These reduced datasets were then used as the input features for training our regression and classification models.

#### NCIP series models

For the held-out interaction task, interactions were ranked by padj and every other interaction was assigned to test until a test set of 10% of the total data was reached, with the rest of the interactions assigned to the train set. Five-fold cross validation using stratified k-fold splits was used to explore different L2 weights, with oversampling of high interactions so that those with an LFC > 5 make up the same fraction of the dataset as those below 5. L2 weights were selected from the range 10^−5^ to 10^5^. For every model and input set examined, a model with the same L2 range was trained using the padj values for the training set as training weights to see if incorporating information quality improved model performance.

For the held-out protein task, all 12 proteins were used as the test protein for one set of models, and all six possible designed pairs were used as test sets (P1&P2, P3&P4, …, P11&P12). The other interactions were assigned to the train set. Five-fold cross-validation was used to explore different L2 weights again. L2 weights were selected from the range 10-5 to 105.

#### MALb8 models

The modeling process for the mALb proteins was the same as the NCIP proteins - however, the held out interaction task used test set sizes of 10% of the total dataset, 43 PPIs, 84 PPIs, and 123 PPIs. For the held out protein task, there were no clear pairs or groups of proteins to sample for the test sets. Instead, one, two, or three proteins were randomly drawn from the total set of proteins to make up a test set. Twelve sets of one, two, or three proteins were sampled from the mALB set for test sets to understand training set size effects on model performance.

### Structure visualization

#### Visualization of 3D structures

All structural images were generated using PyMOL version 2.5.2.

#### Structure prediction

For non-DHD proteins without PDB codes noted, structures for visualization purposes were generated with ESMFold (Lin et al., 2022).

## Supplementary Figures

**Figure S1.**
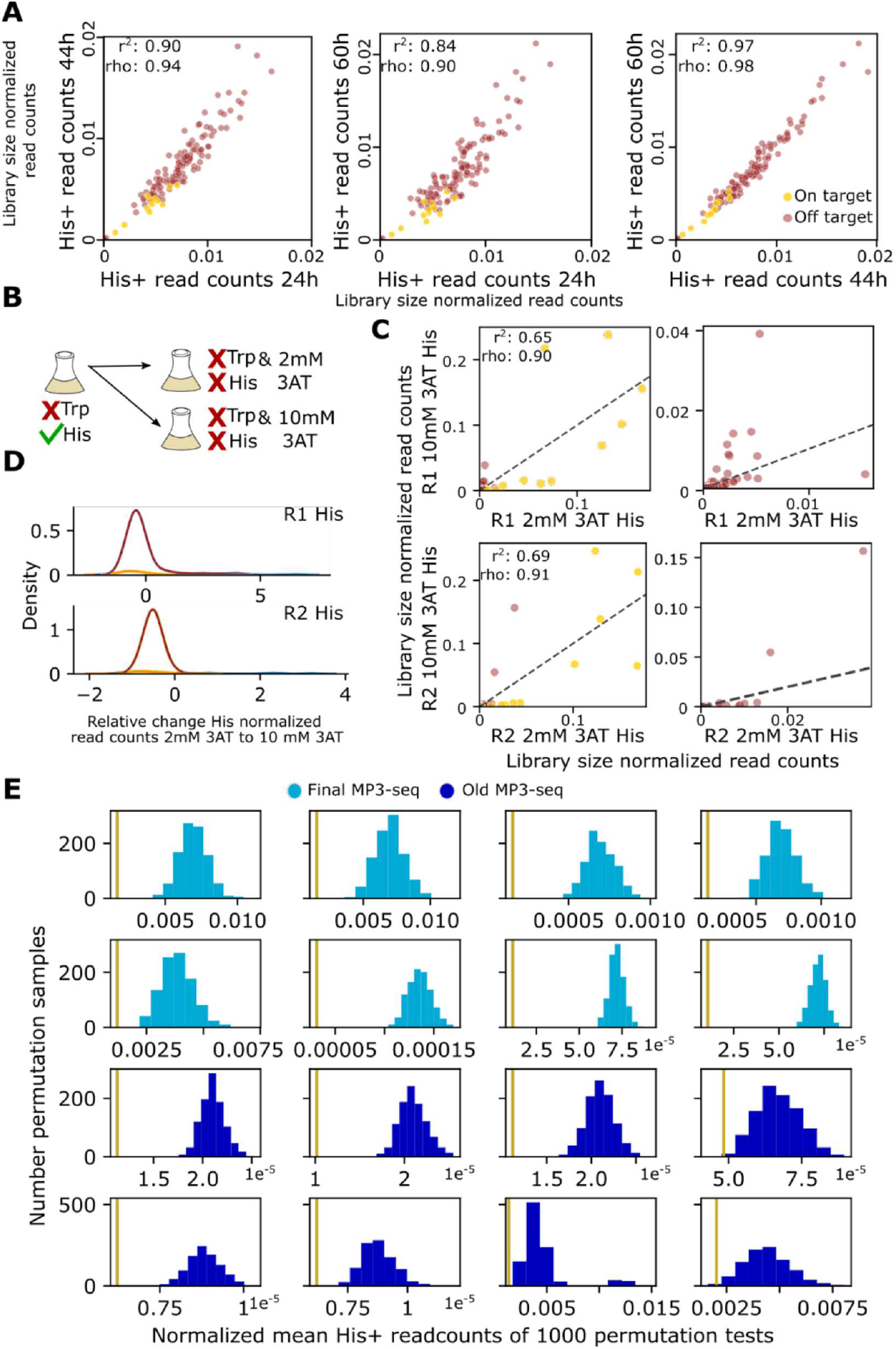
Confirming pre-His and post-His selections proceeded as expected. **A**. Pre-His selection barcode counts hold steady over three samples taken from a P-series all by all screen at 24, 44, and 60 h. **B**. Two concentrations of 3-AT were tested during selection for two P-series all-by-all replicates. **C**. Higher concentration 3-AT post-His selection barcode counts are lower when normalized to library size compared to lower 3-AT selection, as is expected. **D**. Relative changes between the two 3-AT conditions for both replicates. **E**. Permutation test of library size normalized His+ readcounts per library screened, where random samples equal to the number of homodimers in the library are sampled and averaged n=1000 times. All libraries have p<0.01 of the homodimer average being less than heterodimer averages.

**Figure S2.**
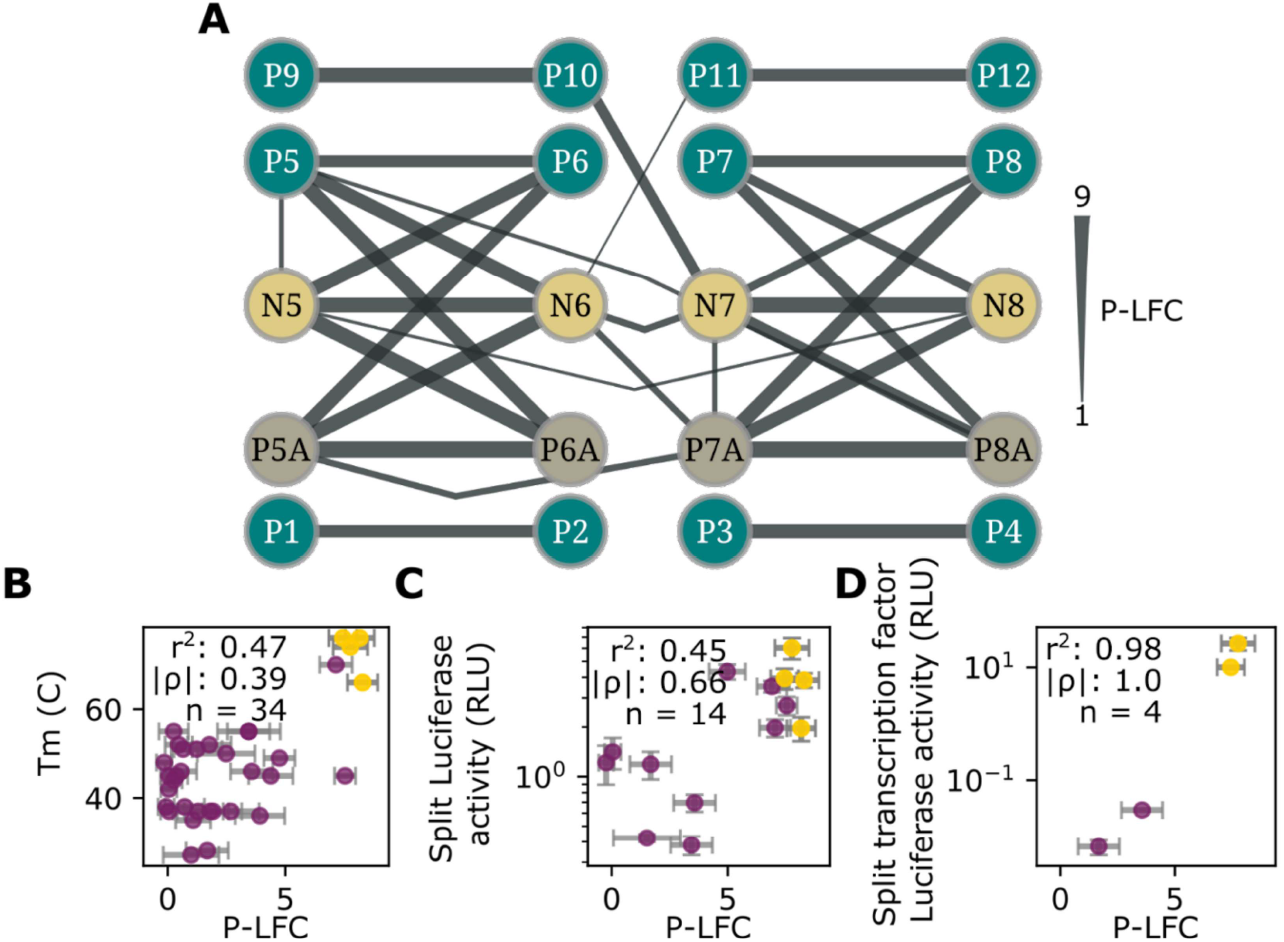
P, PA, and N series designed coil interaction benchmarking and full network. **A** The full undirected P, PA, and N series designed coil interactions from (Plaper et al., 2021) with P_adj_ ≤ 0.05. Line weights correspond to MP3-Seq P-LFCs. **B**. Correlation of P-LFC with calculated midpoint melting temperatures (Tm) from thermal denaturation scans by (Plaper et al., 2021). **C**. Correlation of P-LFCs with split luciferase reconstitution activity in HEK293T cells from (Plaper et al., 2021) (nluc 30 ng and clue 30 ng values used from the split luciferase assays). **D**. Correlation P-LFCs with the split transcription factor assay from (Plaper et al., 2021) (using the 30 ng VP16 and 30 ng TALE-A data).

**Figure S3.**
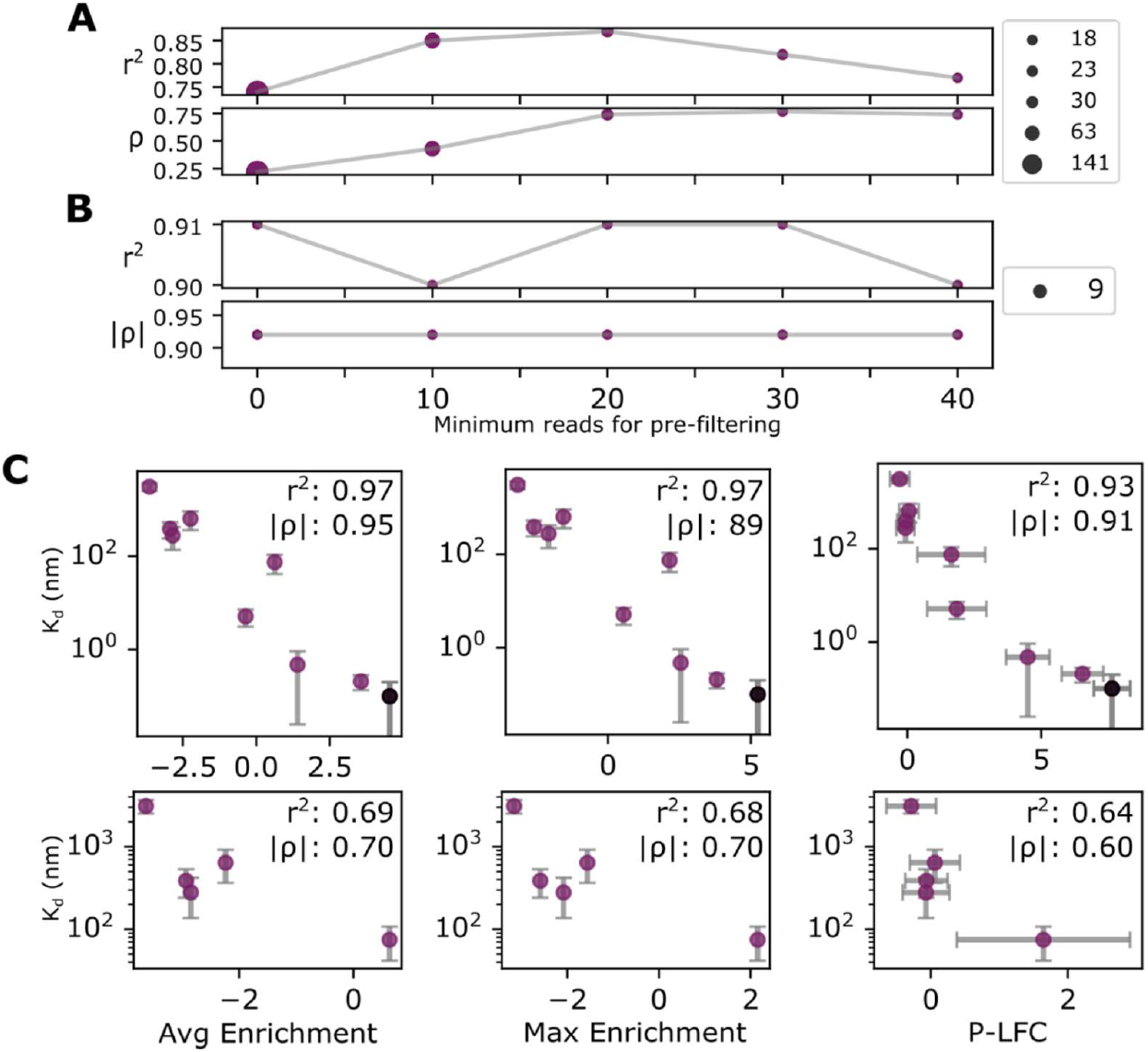
Comparing LFC, PLFC, and enrichment correlations with two benchmark sets. Correlations of P-LFC with the NICP-series fold activation **(A)** and AN/BN binder Kds **(B)** using different minimum barcode count thresholds to pre-filter the PPIs before DESeq2 processing. Sizes of the resulting reduced datasets are shown by the size of the points. C. Correlations of the max and average of both fusion version enrichments per PPI with the Kd values from (Thomas et al., 2013) for all data points (top) and the subset of the weakest interactions (bottom). **C**. The correlation of all points (top) and the weakest interactions (bottom) for average enrichment per PPI across all three replicates, max enrichment per PPI across all three replicates, and the P-LFC values with the Kd values from (Thomas et al., 2013).

**Figure S4.**
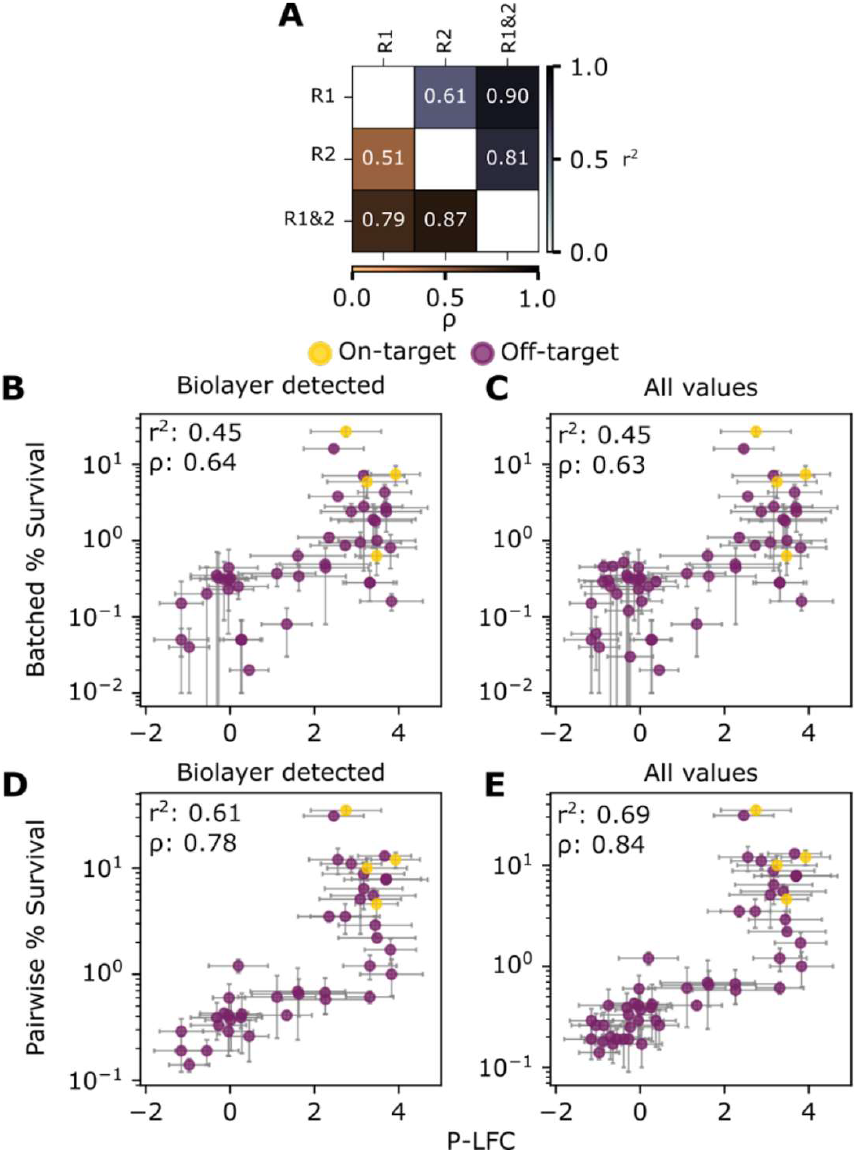
Alpha-Seq correlations with MP3-seq. **A**. Correlation between earlier assay version replicate (R1), final assay version replicate (R2), and DESeq2 combined replicates P-LFC values for the BCL-2 homolog and inhibitor interactions. **B**. MP3-Seq P-LFC versus batched survival for only interactions which succeeded biolayer interferometry C. MP3-Seq P-LFC versus batched survival for all BCL-2 homolog and inhibitor interactions **D**. MP3-Seq P-LFC versus pairwise survival for only interactions which succeeded biolayer interferometry **E**. MP3-Seq P-LFC versus pairwise survival for all BCL-2 homolog and inhibitor interactions

**Figure S5.**
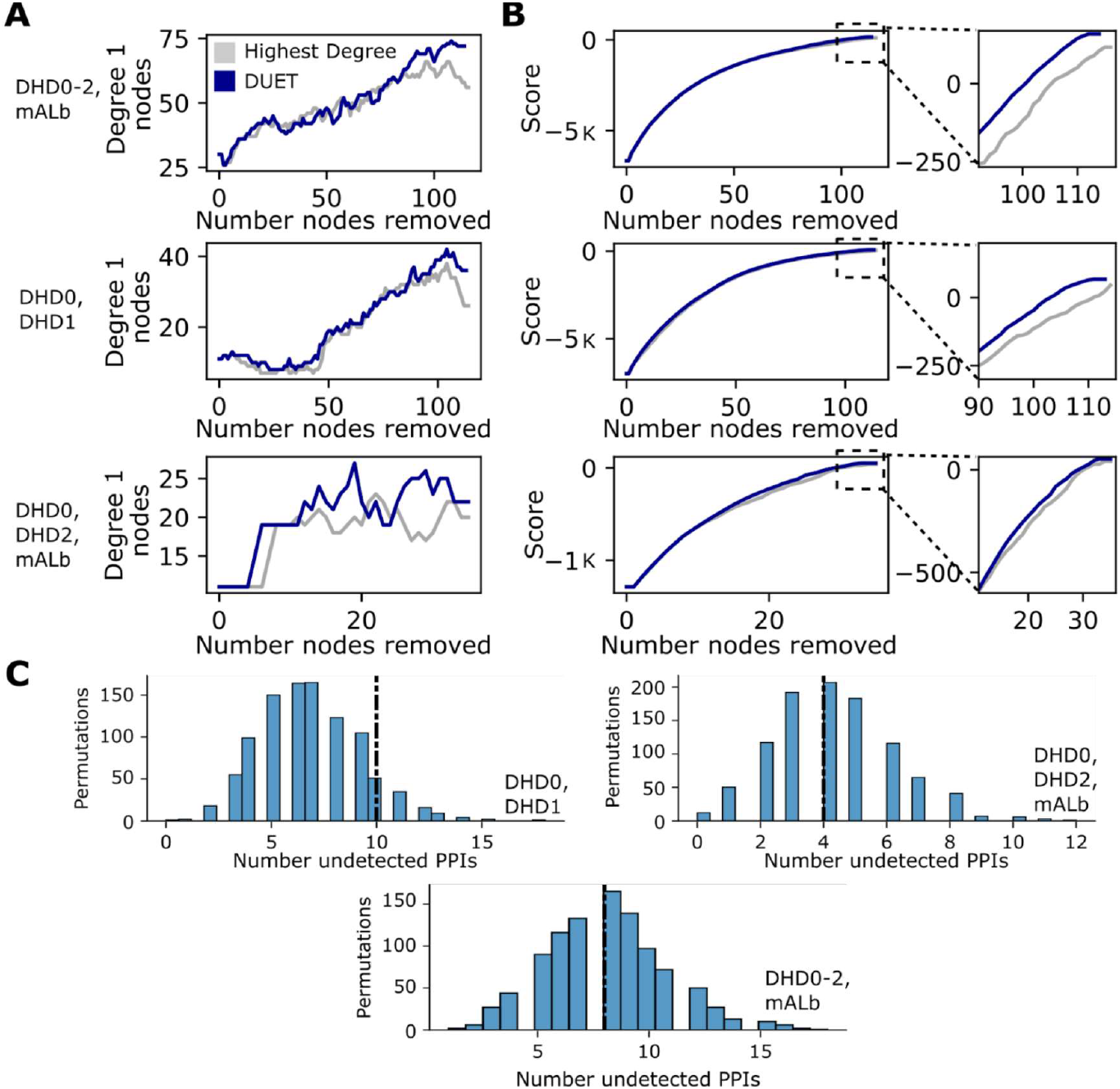
DUET progress per set and permutation tests. **A**. Number of degree one vertices in the three graphs per timestep of the greedy algorithm. For comparison, a similar graph reduction algorithm where the highest degree vertex is removed at every timestep is shown. **B**. Score of the graphs DUET was run on per number of nodes removed, with enlarged portions showing the difference between DUET using the score function and removal of high degree vertices in the final steps. **C**. The number of non-detected interactions in the DUET solution (dashed lines) versus 1000 randomly sampled subsets with the same number of proteins as the DUET solution. Note that the DUET solutions do not contain more PPIs undetected by MP3-Seq in the assay than other proteins.

**Figure S6.**
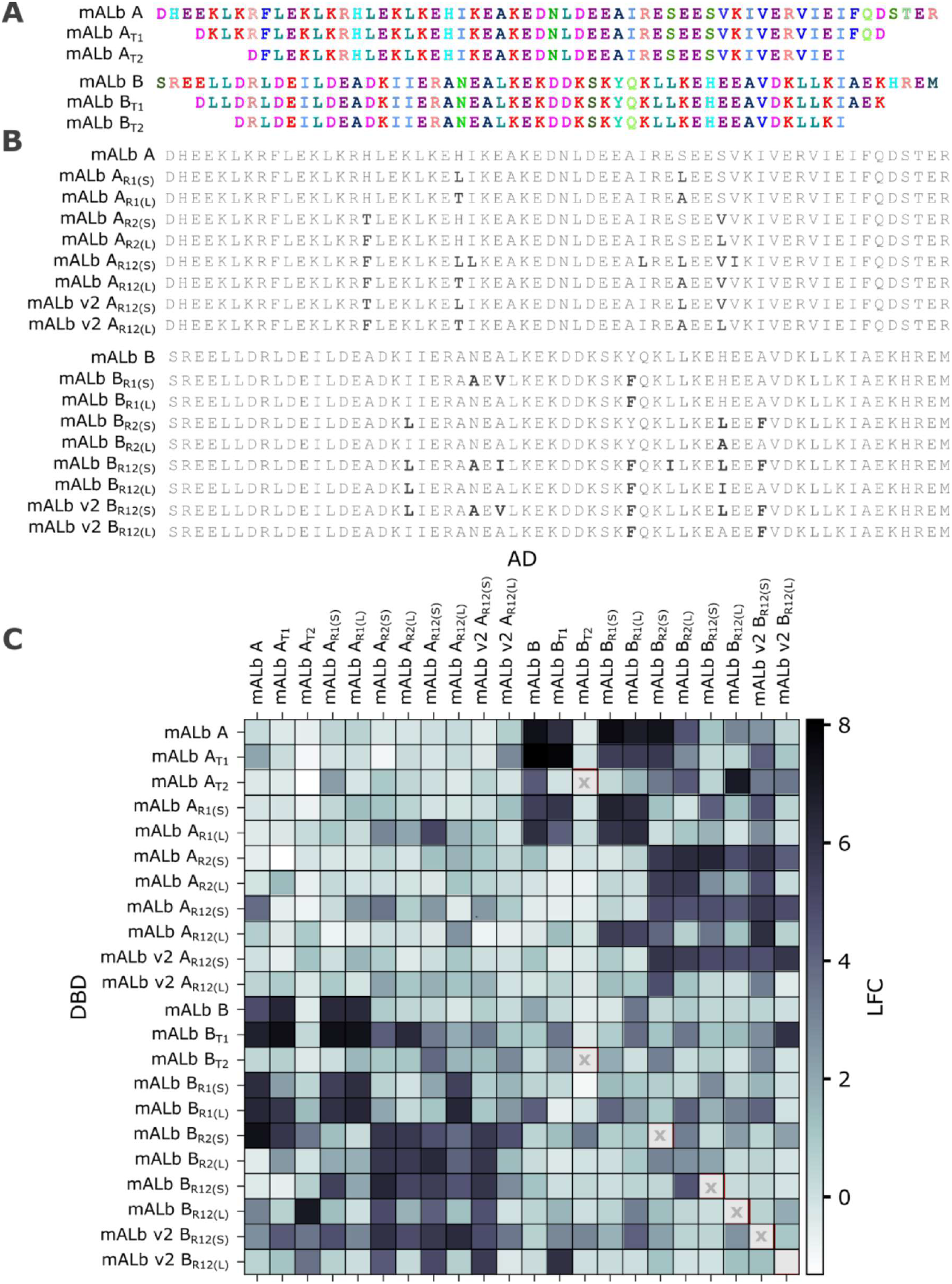
Alignments for mALb8 variant sequences and full interaction heatmap. **A**. mALb truncation sequences **B**. mAlb HB-Net removal sequences, mutated positions are bolded **C**. Heatmap of all mALb variant interaction LFCs.

**Figure S7.**
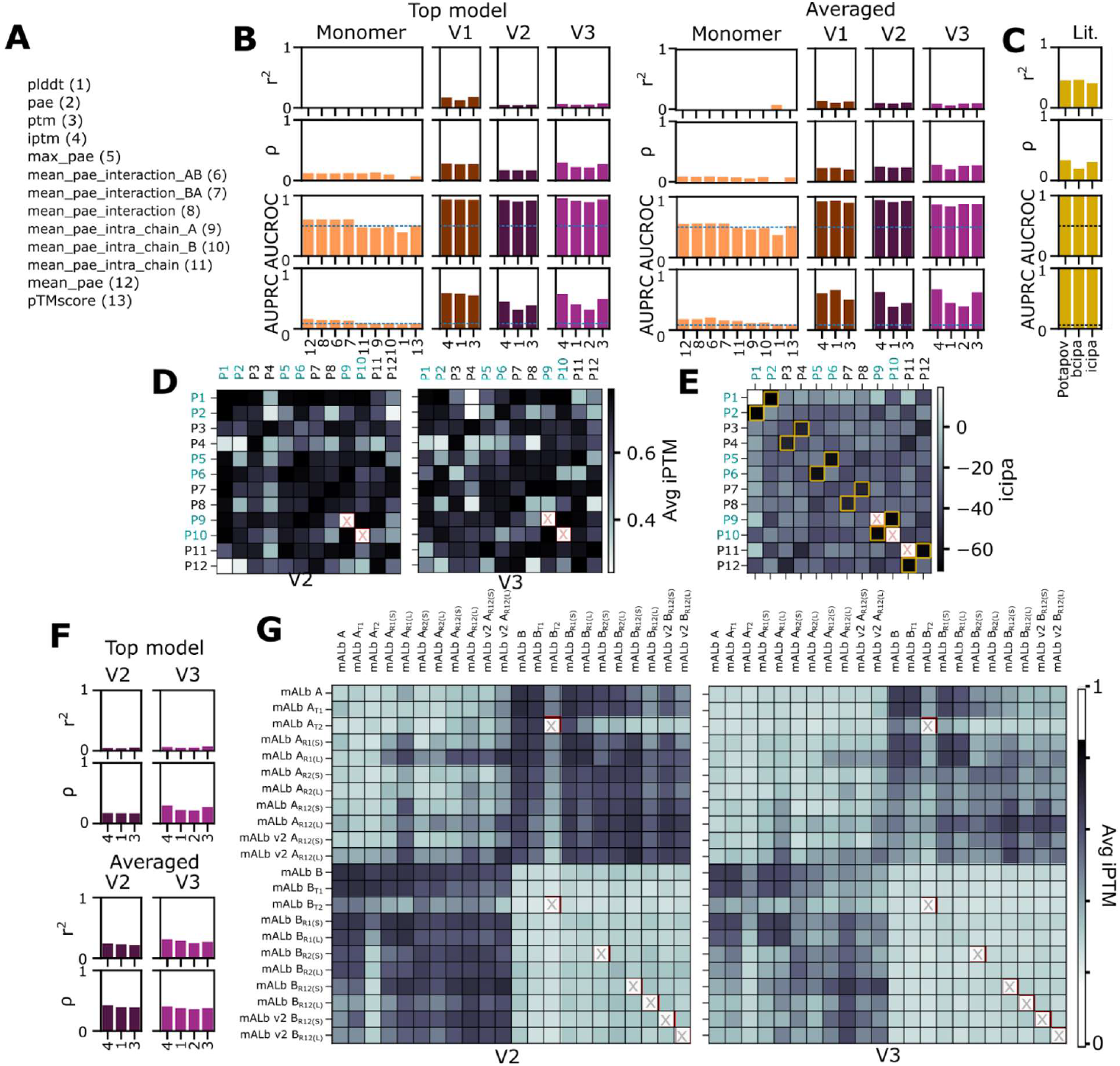
AlphaFold error metrics for the NICP and mALb interactions. **A**. Error metrics examined **B**. Correlation and classification of PNIC LFCs and interactions for top ranked models and averaged models. **C**. Correlation and classification of PNIC interactions using 1×1 predictors (Potapov: (Potapov et al., 2015), bCIPA: (Mason et al., 2006)semi, iCIPA :(Boldridge et al., 2020)). **D**. Average iPTM for the NICP series. **E**. iCipa (1) values for NICP series. **F**. Correlation of AF-M metrics with mALb series LFCs for the top rank model and averages of five models. G. Average iPTM values for the mALb interactions.

**Figure S8.**
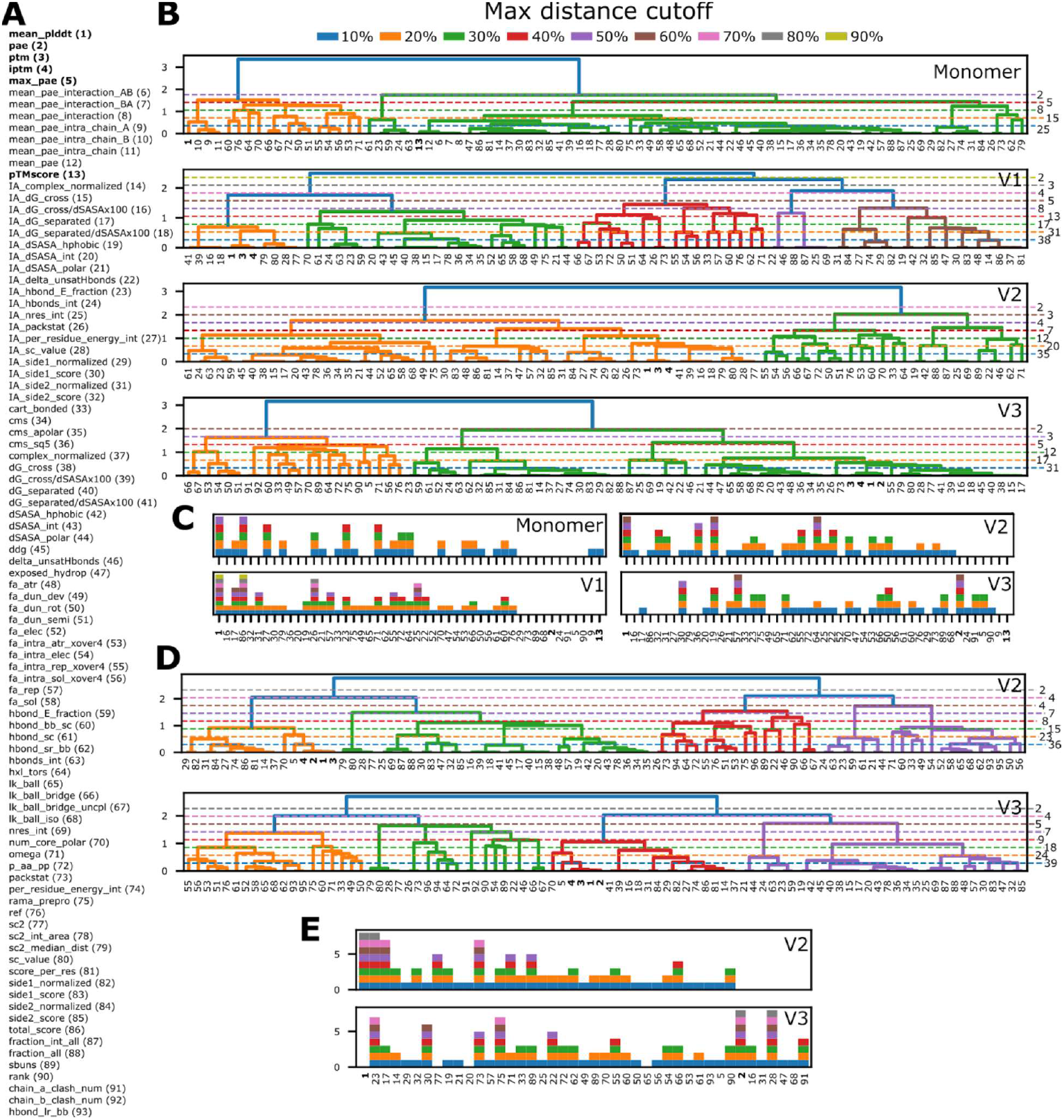
Co-linear feature reduction and list of all features. **A**. All features across models with constant columns removed, bolded features correspond to V1-V3 error metrics. **B**. Feature collinearity reduction for the P-series feature sets across AlphaFold2 (Monomer), AF-M v1, AF-M v2, and AF-M v3. **C**. Features selected per cutoff for the P-series sets. **D**. Feature collinearity reduction for the mALb8 set for AF-M v2 and AF-M v3. **E**. Features selected per cutoff for the mALb8 sets.

**Figure S9.**
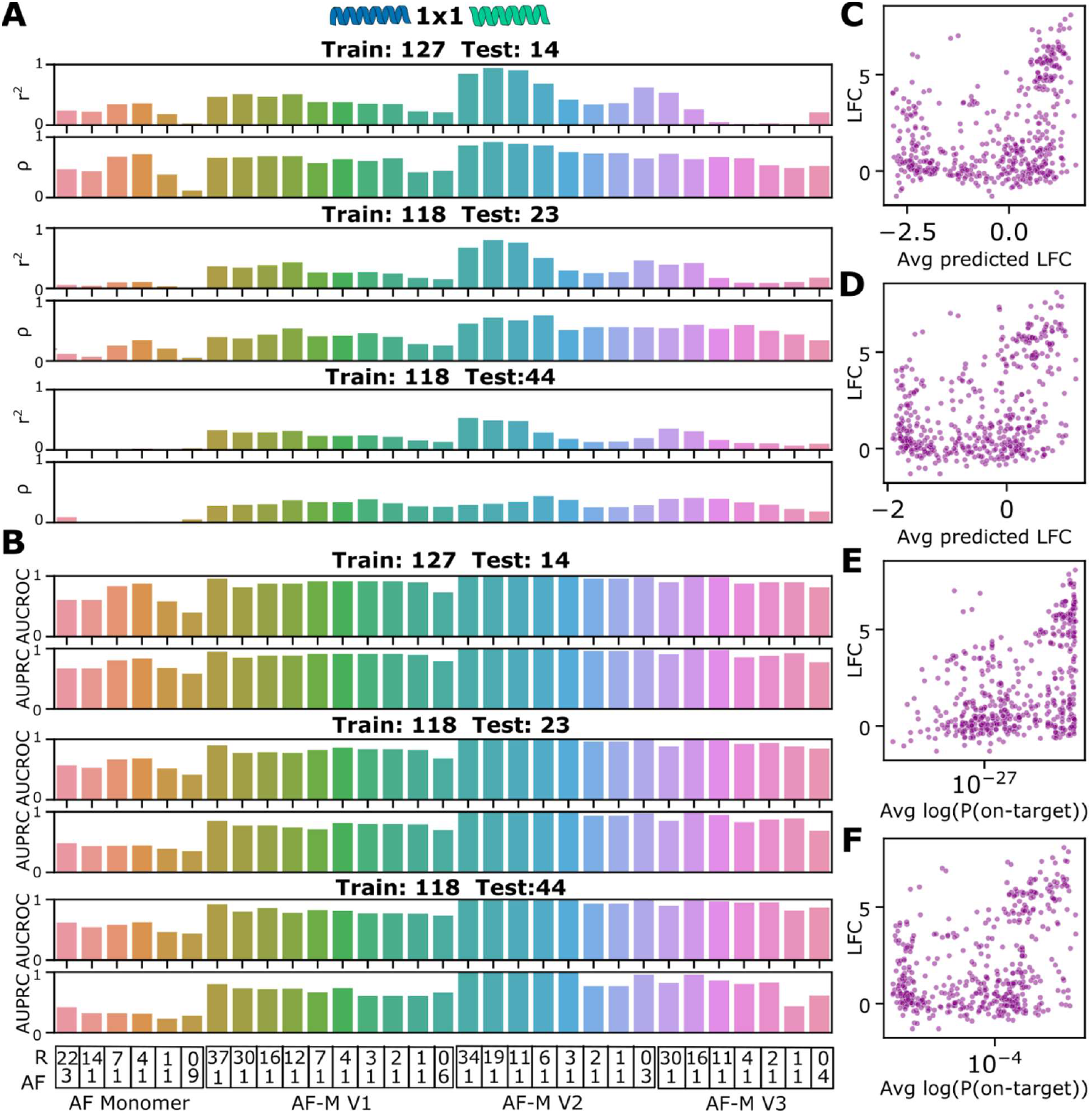
Performance of NICP LFC predictors and classifiers. **A**. Results for heldout interaction task Ridge regression models using varying sizes of train/test sets. The top is a 90/10 train/test split, while the bottom two are for comparison with the held out protein models **B**. Results for the heldout interaction task logistic regression classifiers. The top is a 90/10 train/test split, while the bottom two are for comparison with the held out protein models Number of Rosetta (R) and AF features used as inputs for each model shown. The same number of input features are used for both the regression and classification models. Prediction of mALb interactions with **(C)** the AF feature only AF-M v2 NCIP model and **(D)** the AF feature only AF-M v3 NCIP model. The log probabilities of being “on-target” for the mALb interaction predicted with the NCIP v2 **(E)** and v2 **(F)** AF-M NCIP classifiers.

**Figure S10.**
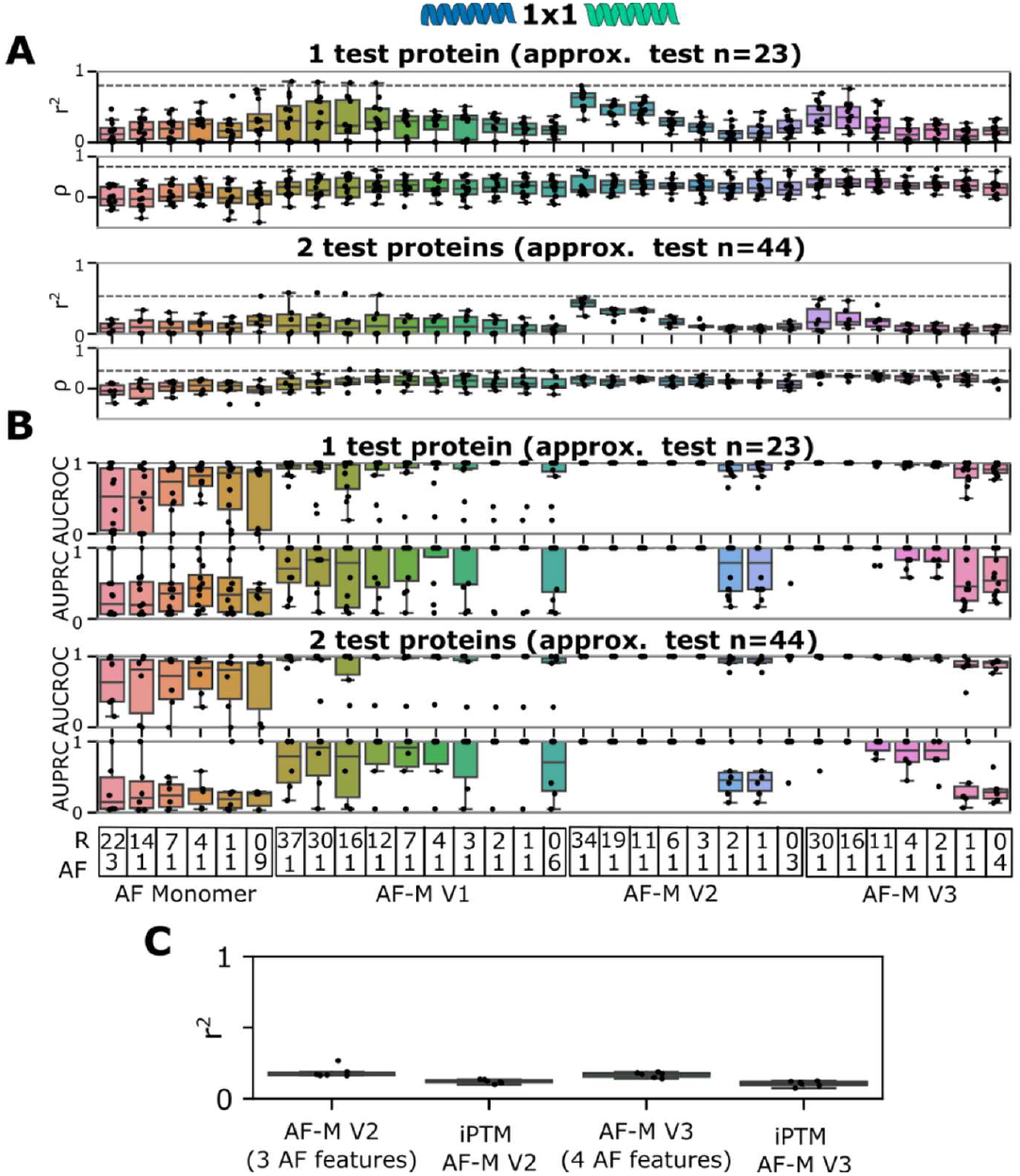
Performance of NICP LFC predictors and classifiers on held out protein sets. **A**. Results for heldout protein task Ridge regression models holding out one and two proteins. The dotted lines correspond to the best-performing held out interaction model with a similarly-sized train/test split. **B**. Results for the heldout interaction task logistic regression classifiers holding out one and two proteins. Number of Rosetta (R) and AFeatures used as inputs for each model shown. **C**. Performance on the training set for holding out two proteins for the regression NCIP models using only AF features.

**Figure S11.**
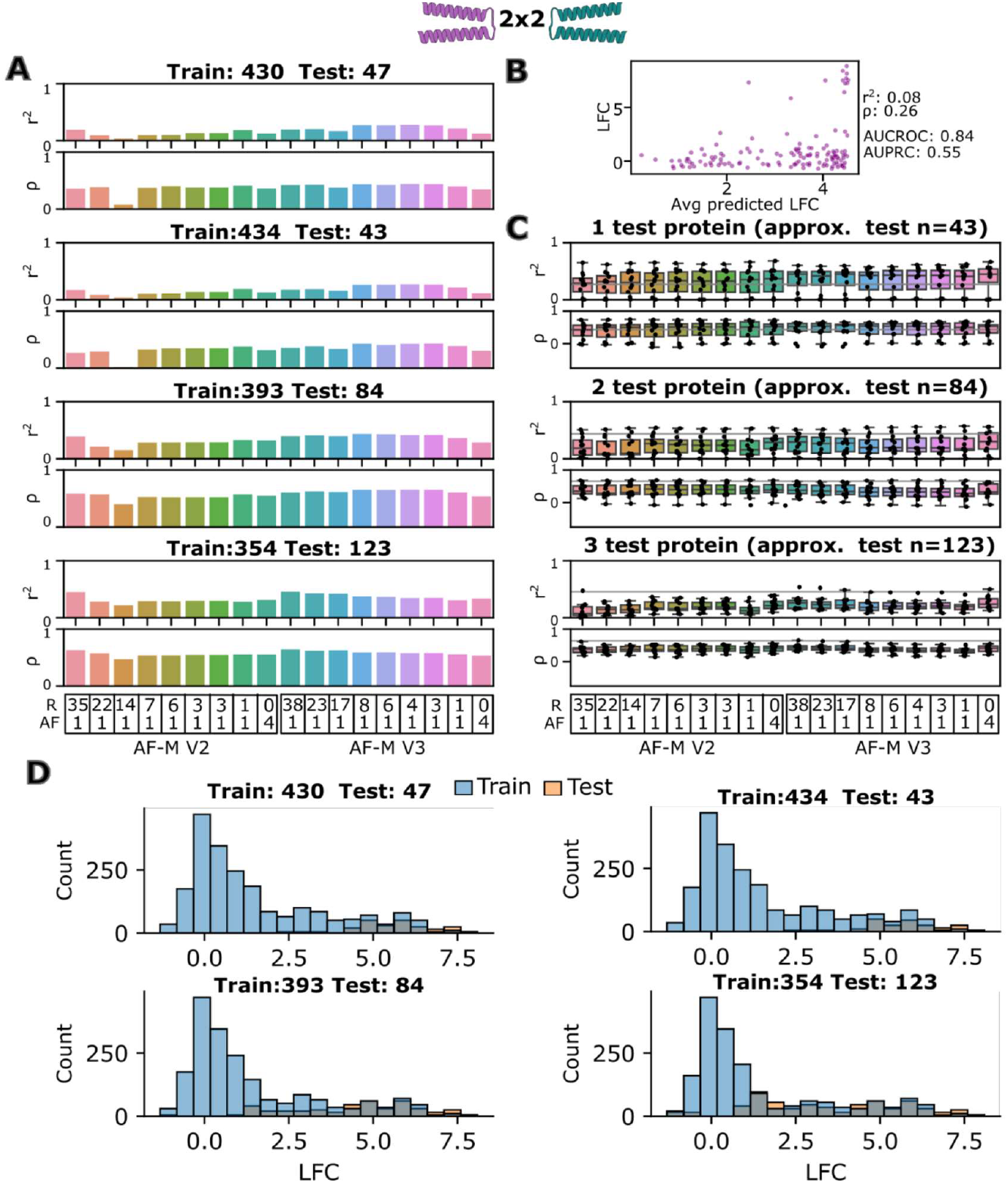
Performance of mALb LFC predictors and prediction of NCIP interactions with mALB models. **A**. Results for heldout interaction task Ridge regression models using varying sizes of train/test sets. The top is a 90/10 train/test split, while the bottom two are for comparison with the held out protein models **B**. Results for heldout protein task Ridge regression models holding out one, two, and three proteins. The dotted lines correspond to the best-performing held out interaction model with a similarly-sized train/test split. Number of Rosetta (R) and AF features used as inputs for each model shown. The same number of input features are used for both the regression and classification models. **C**. Prediction of NCIP interactions with the AF feature only AF-M v2 mALb model. D. Distributions of train and test sets for held-out interaction models.

**Table S1.** MP3-Seq versions, screen sizes, 3-AT concentrations, and autoactivators.

| Experiment | MP3-Seq version | No. proteins | Protein sets used | 3-AT | Autoactivators |
| --- | --- | --- | --- | --- | --- |
| L39 (1) | V1 | 218 | DHD0, DHD1 | 0 | A_HT_DHD_100_swap, A_HT_DHD_33_swap, A_HT_DHD_96_swap, A_HT_DHD_97_swap, B_HT_DHD_94_swap |
| L39 (1) | V1 | 218 | DHD0, DHD1 | 0 | A_HT_DHD_97_swap, A_HT_DHD_33_swap |
| L39 (1) | V1 | 218 | DHD0, DHD1 | 0 | A_HT_DHD_100_swap, A_HT_DHD_33_swap, A_HT_DHD_71_swap, A_HT_DHD_96_swap, A_HT_DHD_97_swap, B_HT_DHD_45_swap, B_HT_DHD_94_swap |
| L43 | V1 | 122 | DHD0 (subset), DHD2, mALb (T) | 1 | ZCON_155_cutT5_B |
| L44 | V1 | 337 | DHD0, DHD1, DHD2, mALb (T) | 1 | A_HT_DHD_33_swap, A_HT_DHD_97_swap, B_HT_DHD_29_swap, Bcl-B, mALb8_cutT1_B |
| L45 | V1 | 15 | Bcl & inhibitors | 1 |  |
| L49 | V1 | 337 | DHD0, DHD1, DHD2, mALb (T) | 1 |  |
| L61 | V2 | 12 | NICP | 2 |  |
| L61 | V2 | 12 | NICP | 10 |  |
| L62 | V2 | 12 | NICP | 2 |  |
| L62 | V2 | 12 | NICP | 10 |  |
| L66 | V2 | 15 | Bcl & inhibitors | 1 |  |
| L67 | V2 | 38 | NICP, mALB (T) | 1 | IL14_A, Jerala_P11 |
| L68 | V2 | 38 | An/Bn, NICP, N, PA series | 1 | AN4, Jerala_P11, N1 |
| L70 | V2 | 86 | mALb (R) | 1 | 1003_mALb8x2_fdrtc_B, 1004_mALb8x2_rprtc_B, 1005_mALb8x12_fdrtc_B, 1007_mALb8x12j_fdrtc_B, 1008_mALb8x12j_rprtc_B, Jerala_P11 |
| L71 | V2 | 118 | NICP, mALB (T), mALb (R), An/Bn, P, N, PA series | 1 | 1004_mALb8x2_rprtc_B, 1005_mALb8x12_fdrtc_B, 1007_mALb8x12j_fdrtc_B, 1008_mALb8x12j_rprtc_B, AN4, IL14_A, MCL1, N1 |
| L72 | V2 | 118 | NICP, mALB (T), mALb (R), An/Bn, P, N, PA series | 1 | 1004_mALb8x2_rprtc_B, 1005_mALb8x12_fdrtc_B, 1007_mALb8x12j_fdrtc_B, 1008_mALb8x12j_rprtc_B, AN4, N1 |

## Bibliography

Baek, M., DiMaio, F., Anishchenko, I., Dauparas, J., Ovchinnikov, S., Lee, G.R., Wang, J., Cong, Q., Kinch, L.N., Schaeffer, R.D., Millan, C., Park, H., Adams, C., Glassman, C.R., DeGiovanni, A., Pereira, J.H., Rodrigues, A.V., van Dijk, A.A., Ebrecht, A.C., Opperman, D.J., Sagmeister, T., Buhlheller, C., Pavkov-Keller, T., Rathinaswamy, M.K., Dalwadi, U., Yip, C.K., Burke, J.E., Garcia, K.C., Grishin, N.V., Adams, P.O., Read, R.J., Baker, D., 2021. Accurate prediction of protein structures and interactions using a three-track neural network. Science 373, 871–876. https://doi.org/10.1126/science.abj8754

Banerjee, S., Velasquez-Zapata, V., Fuerst, G., Elmore, J.M., Wise, R.P., 2021. NGPINT: a next-generation protein-protein interaction software. Brief. Bioinform. 22, bbaa351. https://doi.org/10.1093/bib/bbaa351

Benatuil, L., Perez, J.M., Belk, J., Hsieh, C.-M., 2010. An improved yeast transformation method for the generation of very large human antibody libraries. Protein Eng. Des. Sel. PEDS 23, 155–159. https://doi.org/10.1093/protein/gzq002

Ben-Sasson, A.J., Watson, J.L., Sheffler, W., Johnson, M.C., Bittleston, A., Somasundaram, L., Decarreau, J., Jiao, F., Chen, J., Mela, I., Drabek, A.A., Jarrett, S.M., Blacklow, S.C., Kaminski, C.F., Hura, G.L., De Yoreo, J.J., Kollman, J.M., Ruohola-Baker, H., Derivery, E., Baker, D., 2021. Design of biologically active binary protein 2D materials. Nature 589, 468–473. https://doi.org/10.1038/s41586-020-03120-8

Berger, S., Procko, E., Margineantu, D., Lee, E.F., Shen, B.W., Zelter, A., Silva, D.-A., Chawla, K., Herold, M.J., Garnier, J.-M., Johnson, R., MacCoss, M.J., Lessene, G., Davis, T.N., Stayton, P.S., Stoddard, B.L., Fairlie, W.D., Hackenbery, D.M., Baker, D., 2016. Computationally designed high specificity inhibitors delineate the roles of BCL2 family proteins in cancer. elife 5. https://doi.org/10.7554/elife.20352

Boldridge, W.C., Ljubetič, A., Kim, H., Lubock, N., Szilagyi, D., Lee, J., Brodnik, A., Jerala, R., Kosuri, S., 2020. A Multiplexed Bacterial Two-Hybrid for Rapid Characterization of Protein-Protein Interactions and Iterative Protein Design (preprint). Molecular Biology. https://doi.org/10.1101/2020.11.12.377184

Boyken, S.E., Chen, Z., Groves, B., Langan, R.A., Oberdorfer, G., Ford, A., Gilmore, J.M., Xu, C., DiMaio, F., Pereira, J.H., Sankaran, B., Seelig, G., Zwart, P.H., Baker, D., 2016. De novo design of protein homo-oligomers with modular hydrogen-bond network-mediated specificity. Science 352, 680–687. https://doi.org/10.1126/science.aad8865

Brodnik, A., Jovicic, V., Palangetic, M., Siladi, D., 2019. Construction of orthogonal CC-sets. Informatica 43. https://doi.org/10.31449/inf.v43i1.2693

Chen, Z., Boyken, S.E., Jia, M., Busch, F., Flores-Solis, D., Bick, M.J., Lu, P., VanAernum, Z.L., Sahasrabuddhe, A., Langan, R.A., Bermeo, S., Brunette, T.J., Mulligan, V.K., Carter, L.P., DiMaio, F., Sgourakis, N.G., Wysocki, V.H., Baker, D., 2019. Programmable design of orthogonal protein heterodimers. Nature 565, 106–111. https://doi.org/10.1038/s41586-018-0802-y

Chen, Z., Elowitz, M.B., 2021. Programmable protein circuit design. Cell 184, 2284–2301. https://doi.org/10.1016/j.cell.2021.03.007

Chen, Z., Kibler, R.D., Hunt, A., Busch, F., Pearl, J., Jia, M., VanAernum, Z.L., Wicky, B.I.M., Dods, G., Liao, H., Wilken, M.S., Ciarlo, C., Green, S., El-Samad, H., Stamatoyannopoulos, J., Wysocki, V.H., Jewett, M.C., Boyken, S.E., Baker, D., 2020. De novo design of protein logic gates. Science 368, 78–84. https://doi.org/10.1126/science.aay2790

Cho, J.H., Collins, J.J., Wong, W.W., 2018. Universal Chimeric Antigen Receptors for Multiplexed and Logical Control of T Cell Responses. Cell 173, 1426–1438.e11. https://doi.org/10.1016/j.cell.2018.03.038

Coventry, B., Baker, O., 2021. Protein sequence optimization with a pairwise decomposable penalty for buried unsatisfied hydrogen bonds. PLOS Comput. Biol. 17, e1008061. https://doi.org/10.1371/journal.pcbi.1008061

Curran, K.A., Morse, N.J., Markham, K.A., Wagman, A.M., Gupta, A., Alper, H.S., 2015. Short Synthetic Terminators for Improved Heterologous Gene Expression in Yeast. ACS Synth. Biol. 4, 824–832. https://doi.org/10.1021/sb5003357

Diss, G., Lehner, B., 2018. The genetic landscape of a physical interaction. elife 7, e32472. https://doi.org/10.7554/elife.32472

Erffelinck, M.-L., Ribeiro, B., Perassolo, M., Pauwels, L., Pollier, J., Storme, V., Goossens, A., 2018. A user-friendly platform for yeast two-hybrid library screening using next generation sequencing. PLOS ONE 13, e0201270. https://doi.org/10.1371/journal.pone.0201270

Fields, S., Song, O., 1989. A novel genetic system to detect protein-protein interactions. Nature 340, 245–246. https://doi.org/10.1038/340245a0

Fleishman, S.J., Leaver-Fay, A., Corn, J.E., Strauch, E.-M., Khare, S.D., Koga, N., Ashworth, J., Murphy, P., Richter, F., Lemmon, G., Meiler, J., Baker, D., 2011. RosettaScripts: A Scripting Language Interface to the Rosetta Macromolecular Modeling Suite. PLOS ONE 6, e20161. https://doi.org/10.1371/journal.pone.0020161

Fong, J.H., Keating, A.E., Singh, M., 2004. Predicting specificity in bZIP coiled-coil interactions. Genome Biol. 5, R11. https://doi.org/10.1186/gb-2004-5-2-r11

Gao, X.J., Chong, L.S., Kim, M.S., Elowitz, M.B., 2018. Programmable protein circuits in living cells. Science 361, 1252–1258. https://doi.org/10.1126/science.aat5062

Gonen, S., DiMaio, F., Gonen, T., Baker, O., 2015. Design of ordered two-dimensional arrays mediated by noncovalent protein-protein interfaces. Science 348, 1365–1368. https://doi.org/10.1126/science.aaa9897

Groves, B., Khakhar, A., Nadel, C.M., Gardner, R.G., Seelig, G., 2016. Rewiring MAP kinases in Saccharomyces cerevisiae to regulate novel targets through ubiquitination. elife 5, e15200. https://doi.org/10.7554/elife.15200

Jin, F., Avramova, L., Huang, J., Hazbun, T., 2007. A yeast two-hybrid smart-pool-array system for protein-interaction mapping. Nat. Methods 4, 405–407. https://doi.org/10.1038/nmeth1042

Johansson-Akhe, I., Wallner, B., 2022. Improving peptide-protein docking with AlphaFold-Multimer using forced sampling. Front. Bioinforma. 2.

Jumper, J., Evans, R., Pritzel, A., Green, T., Figurnov, M., Ronneberger, O., Tunyasuvunakool, K., Bates, R., Zfdek, A., Potapenko, A., Bridgland, A., Meyer, C., Kohl, S.A.A., Ballard, A.J., Cowie, A., Romera-Paredes, B., Nikolov, S., Jain, R., Adler, J., Back, T., Petersen, S., Reiman, D., Clancy, E., Zielinski, M., Steinegger, M., Pacholska, M., Berghammer, T., Bodenstein, S., Silver, D., Vinyals, O., Senior, A.W., Kavukcuoglu, K., Kohli, P., Hassabis, D., 2021. Highly accurate protein structure prediction with AlphaFold. Nature 596, 583–589. https://doi.org/10.1038/s41586-021-03819-2

Kim, I., Miller, C.R., Young, D.L., Fields, S., 2013. High-throughput Analysis of in vivo Protein Stability. Mol. Cell. Proteomics 12, 3370–3378. https://doi.org/10.1074/mcp.O113.031708

Kuriyan, J., Cowburn, D., 1997. Modular peptide recognition domains in eukaryotic signaling. Annu. Rev. Biophys. Biomol. Struct. 26, 259–288. https://doi.org/10.1146/annurev.biophys.26.1.259

Lebar, T., Lainscek, D., Merljak, E., Aupic, J., Jerala, R., 2020. A tunable orthogonal coiled-coil interaction toolbox for engineering mammalian cells. Nat. Chem. Biol. 16, 513–519. https://doi.org/10.1038/s41589-019-0443-y

Lin, Z., Akin, H., Rao, R., Hie, B., Zhu, Z., Lu, W., Smetanin, N., Verkuil, R., Kabeli, O., Shmueli, Y., dos Santos Costa, A., Fazel-Zarandi, M., Sercu, T., Candido, S., Rives, A., 2022. Evolutionary-scale prediction of atomic level protein structure with a language model (preprint). Synthetic Biology. https://doi.org/10.1101/2022.07.20.500902

Ljubetič, A., Gradisar, H., Jerala, R., 2017a. Advances in design of protein folds and assemblies. Curr. Opin. Chem. Biol. 40, 65–71. https://doi.org/10.1016/j.cbpa.2017.06.020

Ljubetič, A., Lapenta, F., Gradisar, H., Drobnak, I., Aupic, J., Strmsek, Lainscek D., Hafner-Bratkovic, I., Majerle, A., Krivec, N., Bencina, M., Pisanski, T., Velickovic, T.C., Round, A., Carazo, J.M., Melero, R., Jerala, R., 2017b. Design of coiled-coil protein-origami cages that self-assemble in vitro and in vivo. Nat. Biotechnol. 35, 1094–1101. https://doi.org/10.1038/nbt.3994

Love, M.I., Huber, W., Anders, S., 2014. Moderated estimation of fold change and dispersion for RNA-seq data with DESeq2. Genome Biol. 15, 550. https://doi.org/10.1186/s13059-014-0550-8

Luck, K., Kim, D.-K., Lambourne, L., Spirohn, K., Begg, B.E., Bian, W., Brignall, R., Cafarelli, T., Campos-Laborie, F.J., Charloteaux, B., Choi, D., Cote, A.G., Daley, M., Deimling, S., Desbuleux, A., Dricot, A., Gebbia, M., Hardy, M.F., Kishore, N., Knapp, J.J., Kovacs, I.A., Lemmens, I., Mee, M.W., Mellor, J.C., Pollis, C., Pons, C., Richardson, A.O., Schlabach, S., Teeking, B., Yadav, A., Babor, M., Saleha, D., Basha, O., Bowman-Colin, C., Chin, S.-F., Choi, S.G., Colabella, C., Coppin, G., D’Amata, C., De Ridder, D., De Rouck, S., Duran-Frigola, M., Ennajdaoui, H., Goebels, F., Goehring, L., Gopal, A., Haddad, G., Hatchi, E., Helmy, M., Jacob, Y., Kassa, Y., Landini, S., Li, R., van Lieshout, N., MacWilliams, A., Markey, D., Paulson, J.N., Rangarajan, S., Rasla, J., Rayhan, A., Rolland, T., San-Miguel, A., Shen, Y., Sheykhkarimli, D., Sheynkman, G.M., Simonovsky, E., Ta an, M., Tejeda, A., Tropepe, V., Twizere, J.-C., Wang, Y., Weatheritt, R.J., Weile, J., Xia, Y., Yang, X., Yeger-Lotem, E., Zhong, Q., Aloy, P., Bader, G.D., De Las Rivas, J., Gaudet, S., Hao, T., Rak, J., Tavernier, J., Hill, D.E., Vidal, M., Roth, F.P., Calderwood, M.A., 2020. A reference map of the human binary protein interactome. Nature 580, 402–408. https://doi.org/10.1038/s41586-020-2188-x

Martin, M., 2011. Cutadapt removes adapter sequences from high-throughput sequencing reads. EMBnet.journal 17, 10. https://doi.org/10.14806/ej.17.1.200

Mason, J.M., Schmitz, M.A., Muller, K.M., Arndt, K.M., 2006. Semirational design of Jun-Fos coiled coils with increased affinity: Universal implications for leucine zipper prediction and design. Proc. Natl. Acad. Sci. 103, 8989–8994. https://doi.org/10.1073/pnas.0509880103

Mcisaac, R.S., Oakes, B.L., Wang, X., Dummit, K.A., Botstein, D., Noyes, M.B., 2013. Synthetic gene expression perturbation systems with rapid, tunable, single-gene specificity in yeast. Nucleic Acids Res. 41, e57. https://doi.org/10.1093/nar/gks1313

Mirdita, M., Schutze, K., Moriwaki, Y., Heo, L., Ovchinnikov, S., Steinegger, M., 2022. ColabFold: making protein folding accessible to all. Nat. Methods 19, 679–682. https://doi.org/10.1038/s41592-022-01488-1

Plaper, T., Aupic, J., Dekleva, P., Lapenta, F., Keber, M.M., Jerala, R., Bencina, M., 2021. Coiled-coil heterodimers with increased stability for cellular regulation and sensing SARS-CoV-2 spike protein-mediated cell fusion. Sci. Rep. 11, 9136. https://doi.org/10.1038/s41598-021-88315-3

Potapov, V., Kaplan, J.B., Keating, A.E., 2015. Data-Driven Prediction and Design of bZIP Coiled-Coil Interactions. PLOS Comput. Biol. 11, e1004046. https://doi.org/10.1371/journal.pcbi.1004046

Rajagopala, S.V., Uetz, P., 2009. Analysis of Protein-Protein Interactions Using Array-Based Yeast Two-Hybrid Screens, in: Stagljar, I. (Ed.), Yeast Functional Genomics and Proteomics, Methods in Molecular Biology. Humana Press, Totowa, NJ, pp. 223–245. https://doi.org/10.1007/978-1-59745-540-4_13

Rao, V.S., Srinivas, K., Sujini, G.N., Kumar, G.N.S., 2014. Protein-Protein Interaction Detection: Methods and Analysis. Int. J. Proteomics 2014, 1–12. https://doi.org/10.1155/2014/147648

Rocklin, G.J., Chidyausiku, T.M., Goreshnik, I., Ford, A., Houliston, S., Lemak, A., Carter, L., Ravichandran, R., Mulligan, V.K., Chevalier, A., Arrowsmith, C.H., Baker, D., 2017. Global analysis of protein folding using massively parallel design, synthesis, and testing. Science 357, 168–175. https://doi.org/10.1126/science.aan0693

Rogers, J.M., Wong, C.T., Clarke, J., 2014. Coupled Folding and Binding of the Disordered Protein PUMA Does Not Require Particular Residual Structure. J. Am. Chem. Soc. 136, 5197–5200. https://doi.org/10.1021/ja4125065

Rouet, R., Jackson, K.J.L., Langley, D.B., Christ, D., 2018. Next-Generation Sequencing of Antibody Display Repertoires. Front. lmmunol. 9, 118. https://doi.org/10.3389/fimmu.2018.00118

Scanlon, T.C., Gray, E.C., Griswold, K.E., 2009. Quantifying and resolving multiple vector transformants in S. cerevisiae plasmid libraries. BMC Biotechnol. 9, 95. https://doi.org/10.1186/1472-6750-9-95

Shivhare, D., Musialak-Lange, M., Julca, I., Gluza, P., Mutwil, M., 2021. Removing auto-activators from yeast-two-hybrid assays by conditional negative selection. Sci. Rep. 11, 5477. https://doi.org/10.1038/s41598-021-84608-9

Smits, A.H., Vermeulen, M., 2016. Characterizing Protein-Protein Interactions Using Mass Spectrometry: Challenges and Opportunities. Trends Biotechnol. 34, 825–834. https://doi.org/10.1016/j.tibtech.2016.02.014

Thomas, F., Boyle, A.L., Burton, A.J., Woolfson, D.N., 2013. A Set of de Novo Designed Parallel Heterodimeric Coiled Coils with Quantified Dissociation Constants in the Micromolar to Sub-nanomolar Regime. J. Am. Chem. Soc. 135, 5161–5166. https://doi.org/10.1021/ja312310g

Thompson, K.E., Bashor, C.J., Lim, W.A., Keating, A.E., 2012. SYNZIP Protein Interaction Toolbox: in Vitro and in Vivo Specifications of Heterospecific Coiled-Coil Interaction Domains. ACS Synth. Biol. 1, 118–129. https://doi.org/10.1021/sb200015u

Trigg, S.A., Garza, R.M., MacWilliams, A., Nery, J.R., Bartlett, A., Castanon, R., Goubil, A., Feeney, J., O’Malley, R., Huang, S.-S.C., Zhang, Z.Z., Galli, M., Ecker, J.R., 2017. CrY2H-seq: a massively multiplexed assay for deep-coverage interactome mapping. Nat. Methods 14, 819–825. https://doi.org/10.1038/nmeth.4343

Velasquez-Zapata, V., Elmore, J.M., Banerjee, S., Dorman, K.S., Wise, R.P., 2021. Next-generation yeast-two-hybrid analysis with Y2H-SCORES identifies novel interactors of the MLA immune receptor. PLOS Comput. Biol. 17, e1008890. https://doi.org/10.1371/journal.pcbi.1008890

Weile, J., Sun, S., Cote, A.G., Knapp, J., Verby, M., Mellor, J.C., Wu, Y., Pons, C., Wong, C., Lieshout, N., Yang, F., Tasan, M., Tan, G., Yang, S., Fowler, D.M., Nussbaum, R., Bloom, J.D., Vidal, M., Hill, D.E., Aloy, P., Roth, F.P., 2017. A framework for exhaustively mapping functional missense variants. Mol. Syst. Biol. 13, 957. https://doi.org/10.15252/msb.20177908

Weimann, M., Grossmann, A., Woodsmith, J., Ozkan, Z., Birth, P., Meierhofer, D., Benlasfer, N., Valovka, T., Timmermann, B., Wanker, E.E., Sauer, S., Stelzl, U., 2013. A Y2H-seq approach defines the human protein methyltransferase interactome. Nat. Methods 10, 339–342. https://doi.org/10.1038/nmeth.2397

Yachie, N., Petsalaki, E., Mellor, J.C., Weile, J., Jacob, Y., Verby, M., Ozturk, S.B., Li, S., Cote, A.G., Mosca, R., Knapp, J.J., Ko, M., Yu, A., Gebbia, M., Sahni, N., Yi, S., Tyagi, T., Sheykhkarimli, D., Roth, J.F., Wong, C., Musa, L., Snider, J., Liu, Y.-C., Yu, H., Braun, P., Stagljar, I., Hao, T., Calderwood, M.A., Pelletier, L., Aloy, P., Hill, D.E., Vidal, M., Roth, F.P., 2016. Pooled-matrix protein interaction screens using Barcode Fusion Genetics. Mol. Syst. Biol. 12, 863. https://doi.org/10.15252/msb.20156660

Yang, F., Lei, Y., Zhou, M., Yao, Q., Han, Y., Wu, X., Zhong, W., Zhu, C., Xu, W., Tao, R., Chen, X., Lin, D., Rahman, K., Tyagi, R., Habib, Z., Xiao, S., Wang, D., Yu, Y., Chen, H., Fu, Z., Cao, G., 2018. Development and application of a recombination-based library versus library high-throughput yeast two-hybrid (RLL-Y2H) screening system. Nucleic Acids Res. 46, e17. https://doi.org/10.1093/nar/gkx1173

Yang, J.-S., Garriga-Canut, M., Link, N., Carolis, C., Broadbent, K., Beltran-Sastre, V., Serrano, L., Maurer, S.P., 2018. rec-YnH enables simultaneous many-by-many detection of direct protein-protein and protein-RNA interactions. Nat. Commun. 9, 3747. https://doi.org/10.1038/s41467-018-06128-x

Younger, D., Berger, S., Baker, D., Klavins, E., 2017. High-throughput characterization of protein-protein interactions by reprogramming yeast mating. Proc. Natl. Acad. Sci. U. S. A. 114, 12166–12171. https://doi.org/10.1073/pnas.1705867114

Yu, D., Chojnowski, G., Rosenthal, M., Kosinski, J., 2022. AlphaPulldown - a Python package for protein-protein interaction screens using AlphaFold-Multimer. https://doi.org/10.1101/2022.08.05.502961

Yu, H., Tardivo, L., Tam, S., Weiner, E., Gebreab, F., Fan, C., Svrzikapa, N., Hirozane-Kishikawa, T., Rietman, E., Yang, X., Sahalie, J., Salehi-Ashtiani, K., Hao, T., Cusick, M.E., Hill, D.E., Roth, F.P., Braun, P., Vidal, M., 2011. Next-generation sequencing to generate interactome datasets. Nat. Methods 8, 478–480. https://doi.org/10.1038/nmeth.1597

Zhao, L., Liu, Z., Levy, S.F., Wu, S., 2018. Bartender: a fast and accurate clustering algorithm to count barcode reads. Bioinformatics 34, 739–747. https://doi.org/10.1093/bioinformatics/btx655

